# The human glucocorticoid receptor variant rs6190 increases blood cholesterol and promotes atherosclerosis

**DOI:** 10.1101/2024.11.27.625727

**Authors:** Hima Bindu Durumutla, April Haller, Greta Noble, Ashok Daniel Prabakaran, Kevin McFarland, Hannah Latimer, Akanksha Rajput, Olukunle Akinborewa, Bahram Namjou-Khales, David Y. Hui, Mattia Quattrocelli

**Affiliations:** Molecular Cardiovascular Biology, Heart Institute, Cincinnati Children’s Hospital Medical Center, Cincinnati, OH, USA; Dept. Pediatrics,University of Cincinnati College of Medicine, Cincinnati, OH, USA; Department of Pathology,University of Cincinnati College of Medicine, Cincinnati, OH, USA; Dept Pharmacology, Physiology and Neurobiology; University of Cincinnati College of Medicine, Cincinnati, OH, USA; Center for Autoimmune Genomics and Etiology, Cincinnati Children’s Hospital Medical Center, Cincinnati, OH, USA

**Keywords:** Rs6190, ER22/23K, glucocorticoid receptor, SNP, cholesterol, atherosclerosis, liver, hiPSCs, hepatocytes, LDL receptor, PCSK9, transactivation

## Abstract

Elevated cholesterol poses cardiovascular risks. The glucocorticoid receptor (GR) harbors a still undefined role in cholesterol regulation. Here, we report that a coding single nucleotide polymorphism (SNP) in the gene encoding the GR, *rs6190*, associated with increased cholesterol in women according to UK Biobank and All Of Us datasets. In SNP-genocopying mice, we found that the SNP enhanced hepatic GR activity to transactivate *Pcsk9* and *Bhlhe40*, negative regulators of low-density lipoprotein (LDL) and high-density lipoprotein (HDL) receptors respectively. In mice, the SNP was sufficient to elevate circulating cholesterol across all lipoprotein fractions and the risk and severity of atherosclerotic lesions on the pro-atherogenic *hAPOE*2/*2* background. The SNP effect on atherosclerosis was blocked by in vivo liver knockdown of *Pcsk9* and *Bhlhe40*. Also, corticosterone and testosterone were protective against the mutant GR program in cholesterol and atherosclerosis in male mice, while the SNP effect was additive to estrogen loss in females. Remarkably, we found that the mutant GR program was conserved in human hepatocyte-like cells using CRISPR-engineered, SNP-genocopying human induced pluripotent stem cells (hiPSCs). Taken together, our study leverages a non-rare human variant to uncover a novel GR-dependent mechanism contributing to atherogenic risk, particularly in women.

**Graphical Abstract:** 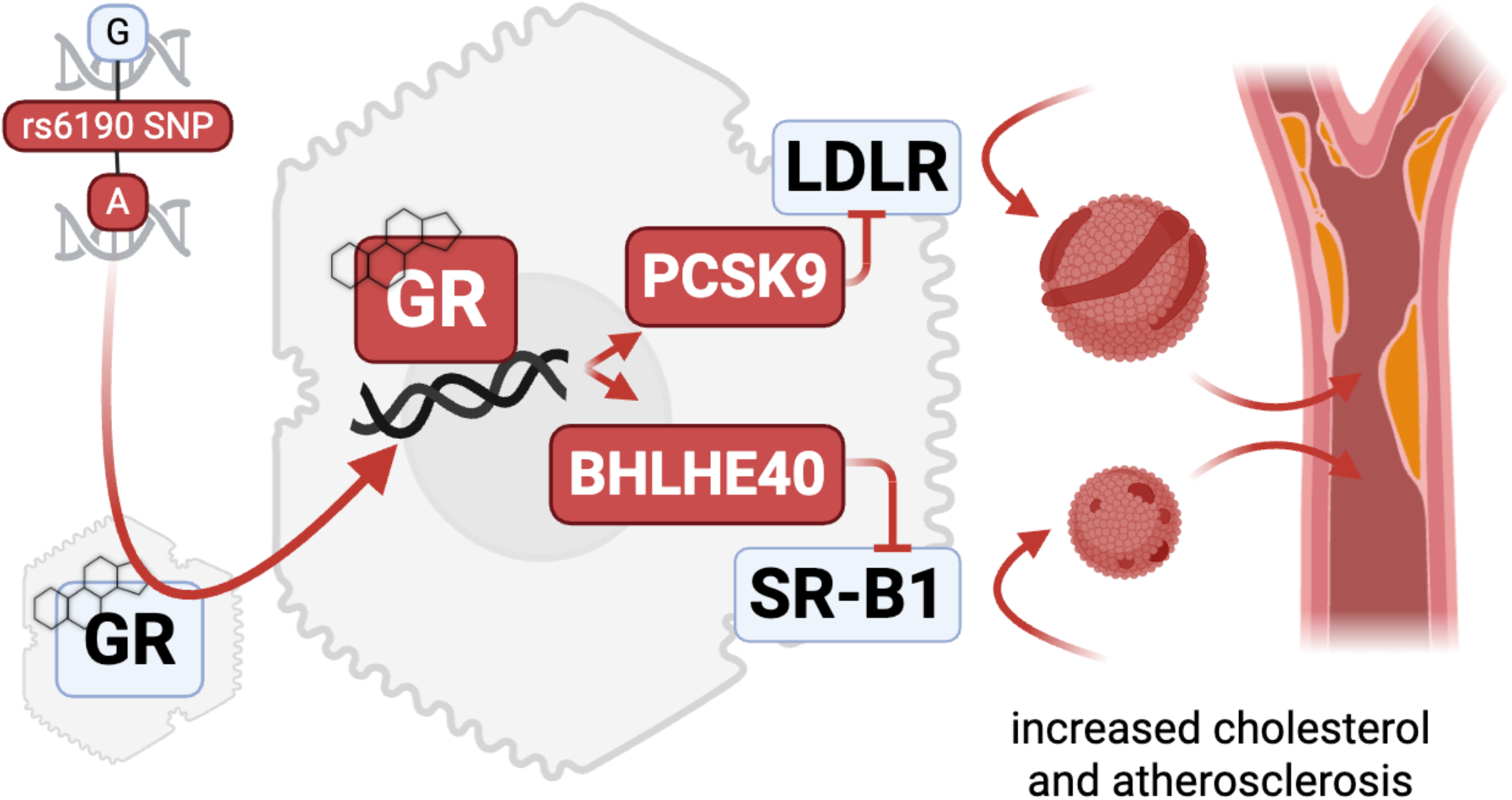

## Introduction

Elevated plasma cholesterol is a significant risk factor for atherosclerotic cardiovascular disease. In women, the risk of developing high cholesterol and atherosclerosis typically increases with age and particularly after meno-pause (1), when the inhibitory effects of estrogen on atherogenesis decline (2). However, the cholesterol regulations underlying this phenomenon remain unclear, as recently shown by counter-intuitive associations between LDL-cholesterol and cardiovascular risk in the UK Biobank (3). Although advancements in drug therapies and lifestyle interventions have demonstrated efficacy, the identification of genetic and epigenetic factors regulating cholesterol is still ongoing to increase our mechanistic understanding and better predict and manage elevated cholesterol.

Despite its involvement in virtually every nutrient metabolism, the glucocorticoid receptor (GR) remains a poorly defined nuclear factor in cholesterol homeostasis. The GR is a ligand-activated nuclear transcription factor that exerts multifaceted effects on nutrient metabolism (4, 5) by transactivating or transrepressing large gene programs in a tissue-specific manner (6). While traditionally recognized for its role in immune regulation, GR pro-foundly influences metabolic processes, including glucose and lipid metabolism (7). Prior studies employing GR knockdown in liver and adipose tissue have shown promising outcomes in mitigating hypercholesterolemia and associated metabolic abnormalities in obese diabetic mice (8). Retrospective studies involving pathomorphological data obtained from human autopsies have provided insights into potential relationships between glucocorticoid treatments and atherogenesis (9-12). However, the direct link between the hepatic GR and regulation of cholesterol levels remains elusive. Indeed, although the glucocorticoid-GR axis has been implicated in apolipoprotein expression (13) and cholesterol efflux in macrophages (14, 15), the epigenetic and transcriptional mechanisms enabled by the GR in hepatocyte-autonomous cholesterol uptake remain still poorly defined.

Previously, several genetic variants in the GR gene (*NR3C1;* OMIM #138040) have been described in the human population. These genetic variants can affect the transcriptional activity of the GR and its downstream target genes, potentially influencing nutrient regulation (16-19). Epidemiological studies have provided evidence of an association between specific GR polymorphisms and variation in lipid profiles (16, 20, 21). Notably, the *rs6190* (c.68G>A; p.R23K) coding single nucleotide polymorphisms (SNP) - also known as “E22R/E23K” due its complete linkage to the silent p.E22E *rs6189* SNP - is a genetic variant in codon 23 that results in an amino acid change from arginine (R) to lysine (K) in the N-terminus of the GR protein (22). This mutation has been linked to alterations in several parameters of metabolic homeostasis in humans, including cholesterol (18). However, the precise molecular mechanisms through which this polymorphism skews GR activity to perturb cholesterol remain poorly characterized.

In this study, we harnessed the human *rs6190* SNP to identify a direct GR-mediated program governing hepatic cholesterol regulation and its association with atherogenic risk. We found that this low-frequency coding SNP correlated with increased levels of cholesterol in women from UK Biobank and All of Us cohorts, and promoted cholesterol and atherosclerosis in transgenic mice according to the number of SNP alleles (homo>hetero>reference). Our transcriptomic and epigenetic datasets revealed that the mutant GR perturbed cholesterol levels through transactivation of *Pcsk9* and *Bhlhe40*, negative regulators of LDL and HDL receptors in the liver and previously unknown targets of GR. Our study identifies rs6190 as a potential risk factor for atherosclerosis, particularly in women, and reports unanticipated mechanisms through which the hepatic GR impacts cholesterol levels in the circulation.

## Results

### rs6190 SNP increases plasma cholesterol levels in women according to allele zygosity

To investigate the influence of rs6190 variant on cholesterol regulation, we probed the large adult cohort from the United Kingdom (UK) Biobank, comprising of 485,895 at the age of 40-70 years. In this cohort, the GR rs6190 variant (*NR3C1* gene, transcript ENST00000231509.3 (-strand); c.68G>A; p.R23K) exhibited a minor allele frequency of 2.75% (25,944 heterozygous, 413 homozygous individuals), categorizing it as a low-frequency variant (22). We screened the quantitated parameters from the NMR metabolomics dataset within the UK Biobank dataset (120,356 individuals comprising of 65156 women and 55380 men; same age range as general dataset, 40-70 years) for rs6190 associations disaggregated by sex. All analyses were adjusted for age, body mass index (BMI), top 10 principal components, and genotype information for 12 commonly-referenced, hypercholesterolemia-associated SNPs within *PCSK9, CELSR2, APOB, ABCG8, SLC22A1, HFE, MYLIP, ST3GAL4, NYNRIN, LDLR*, and *APOE* genes (23). Importantly, none of these 12 classical variants were in the neighborhood of rs6190 and did not show significant pairwise linkage disequilibrium effect (r^2^ < 0.001) at the genomic level. While no associations were significant after multiple testing in men, rs6190 SNP significantly associated with many cholesterol parameters in women, accounting for 23 out of 33 total plasma parameters with a significant rs6190 effect (adjusted p<0.005) (**Figure 1A**).

**Figure 1.**
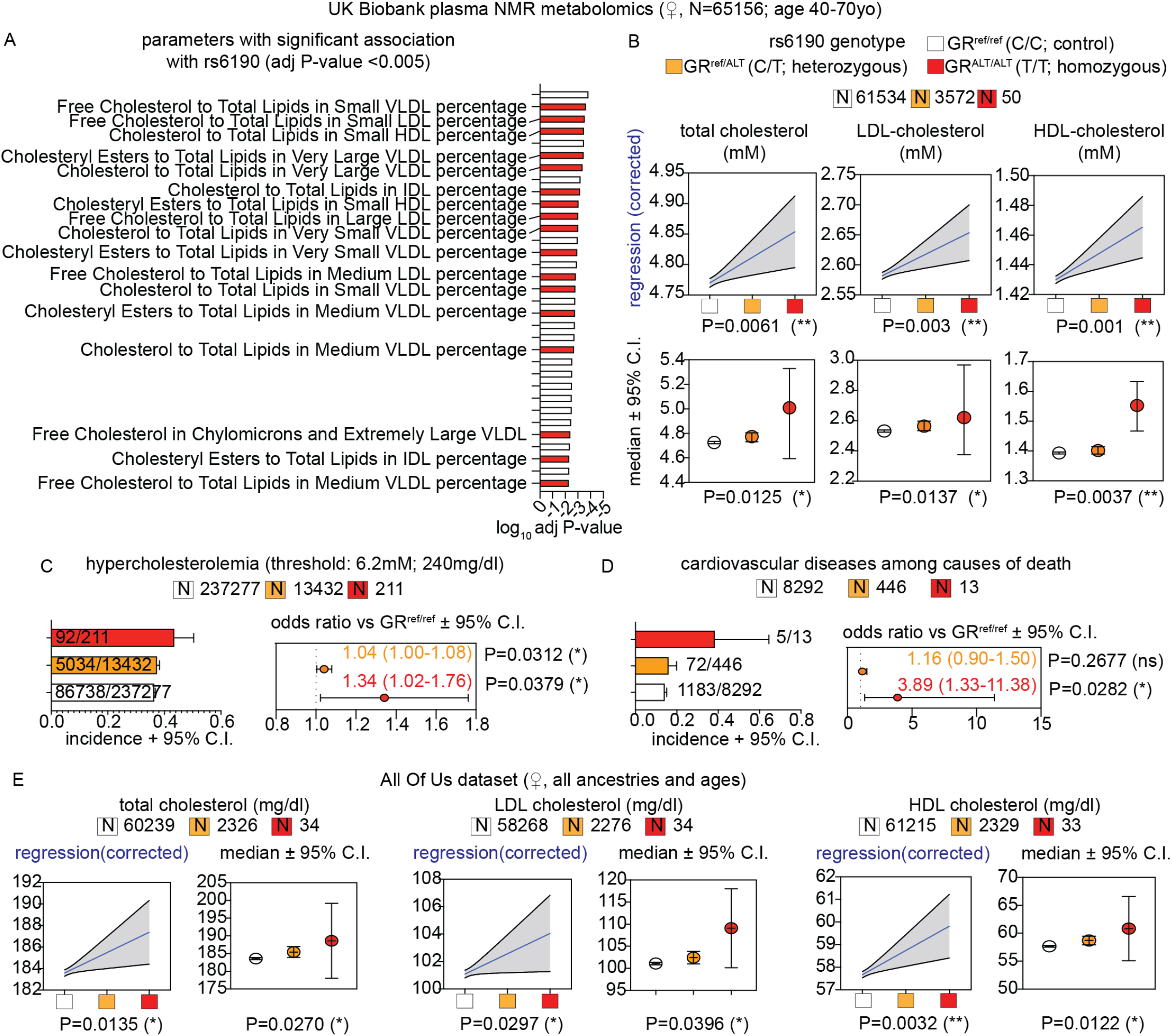
rs6190 correlates with cholesterol increase in women from the UK Biobank and All of US datasets. **(A)** Unbiased ranking of UK Biobank plasma NMR parameters for significant rs6190 effect in women. Cholesterol-related parameters are highlighted in text and red bars. P values were adjusted for age, BMI and canonical hypercholesterolemia-associated SNPs. **(B)** Linear regressions (blue lines; shaded area represents 95% C.I.; corrected for age, diabetes, triacylglycerols) and median confidence intervals (Kruskal-Wallis test) show zygosity-dependent trends in elevation of total, LDL- and HDL-cholesterol in women. **(C-D)** Compared to non-carriers, homozygous SNP carriers showed increased odds ratio for hypercholesterolemia and cardiovascular disease deaths according to ICD10 codes; Chi-square test. **(E)** Linear regressions and median comparisons correlated rs6190 genotype with cholesterol elevation in women from the All of Us dataset, including all ancestries and ages. *, P<0.05; **, P<0.01; ***, P<0.001; ****, P<0.0001.

We then stratified total, LDL-, and HDL-cholesterol values from women according to SNP zygosity. We are defining here homozygous carriers of the reference allele (control population) as GR^ref/ref^, heterozygous SNP carriers as GR^ref/ALT^, and homozygous SNP carriers as GR^ALT/ALT^. We performed linear regressions with a mixed model correcting for age, BMI, diabetic status and triacylglycerols. In parallel, we also compared median confidence intervals across rs6190 genotypes. Remarkably, total, LDL-, and HDL-cholesterol showed a modest but significant elevation of median levels according to the number of SNP alleles in women (**Figure 1B**). The zygosity-dependent trends were not significant in men **(Suppl. Fig. 1A)**. Considering the effects on cholesterol, we probed the total UK Biobank dataset for hypercholesterolemia and cardiovascular disease mortality odds ratios. In alignment with the trends in cholesterol, GR^ALT/ALT^ women displayed an increased odds ratio of 1.34 (95% CI: 1.02 – 1.76; P=0.0092) for hypercholesterolemia (total cholesterol >240 mg/dl) and 2.37 (95% CI: 1.05 – 5.9; P=0.01) for death due to cardiovascular diseases, compared to GR^ref/ref^ women (**Figure 1C-D**).

To probe these rs6190 correlations in a more genetically diverse human dataset, we queried the All Of Us dataset, where we found the SNP at a variable minor allele frequency ranging from low-frequency to rare across ancestries: African/African-American, 0.49%; American Admixed/Latino, 0.84%; East Asian, 0.061%; European, 2.67%; Middle Eastern, 1.43%; South Asian, 1.49%. In the All Of Us subset of 245,385 individuals with rs6190 genotype annotation encompassing all ancestries and ages, we repeated the linear regressions corrected for age, BMI, diabetes, triacylglycerols, as well as the median comparisons. The analyses in the All of Us dataset confirmed a significant correlation between rs6190 zygosity and total, LDL and HDL cholesterol levels in women **(Figure 1E)**, while correlations were not significant once again in men **(Suppl. Fig. 1B)**.

Because *NR3C1* variants did not come up as top hits for cholesterol in GWAS studies so far, we counter-verified whether targeted regressions for total cholesterol could “unmask” potential correlations of *NR3C1* variants not associated with cholesterol through prior GWAS studies. We extracted 38 *NR3C1* locus variants with GWAS significant hits (<5 × 10−8) from the current EBI GWAS catalog and ran linear regressions - aggregated for ancestry but disaggregated for sex - for total cholesterol in the All of Us dataset **(Suppl. Table 1)**. All 38 variants were non-coding and in weak-to-negligible LD range with rs6190 (LD, r^2^<0.15). None of the 38 variants had a significant reported GWAS hit for cholesterol. However, our regressions showed 24 variants with correlations (adj p-val <0.05; 20, direct; 4, inverse) with total cholesterol according to zygosity. Intriguingly, those correlations were significant only in women and not in men for all 24 variants, analogous to our findings with rs6190.

Taken together, our findings highlight the association of the rs6190 SNP with modest but significant elevations of cholesterol in women from the UK Biobank and the All Of Us cohorts.

### The rs6190 SNP is sufficient to increase plasma cholesterol and promotes GR transactivation of Pcsk9 and Bhlhe40 in mice

To elucidate the extent to which the mutant GR promotes cholesterol elevation, we introduced a genocopy of the rs6190 SNP into the endogenous *Nr3c1* (GR gene) locus on the C57BL/6J background. The murine ortholog of the human GR-R23K mutation is GR-R24K due to an additional amino acid in position 10. Employing CRISPR-mediated knock-in recombination, the murine GR gene was targeted at the orthologous codon 24 resulting in C>T mutation in the forward strand (c.71G>A mutation in the codon, reverse strand) leading to a p.R24K amino acid substitution (**Suppl. Figure 2A**). In concordance with human carriers, we define here homozygous mice for wild-type allele as “GR^re/ref^” (control), heterozygous SNP mice as “GR^ref/ALT^”, and homozygous SNP mice as “GR^ALT/ALT^”. In female littermate mice under normal chow conditions, total plasma cholesterol increased according to SNP zygosity in both fasted and fed states (**Figure 2A**). Using the standard fast-performance liquid chromatography (FPLC) method, we found that the GR SNP elicited an increase in cholesterol levels across all lipoprotein fractions – VLDL-, LDL- and HDL-cholesterol - according to SNP allele number, in conditions of either regular chow or 16-week long Western diet in female (**Figure 2B**), but not male mice **(Suppl. Fig. 2B)**. This sex-specific effect in mice paralleled the correlations within human datasets and prompted us to focus on female mice for the bulk of our histological, physiological and mechanistic analyses. After Western diet exposure, in 3 out of 5 analyzed GR^ALT/ALT^ female mice, we found histological evidence of immature plaque formation in the aortic root (**Suppl. Fig. 2C**), a remarkable finding in the absence of pro-atherogenic genetic backgrounds. Our findings provide evidence that, in homogeneous genetic settings, the SNP is sufficient to modestly but significantly elevate total, LDL-, and HDL-cholesterol in females according to an incomplete dominance model, i.e. commensurate to SNP zygosity.

**Figure 2.**
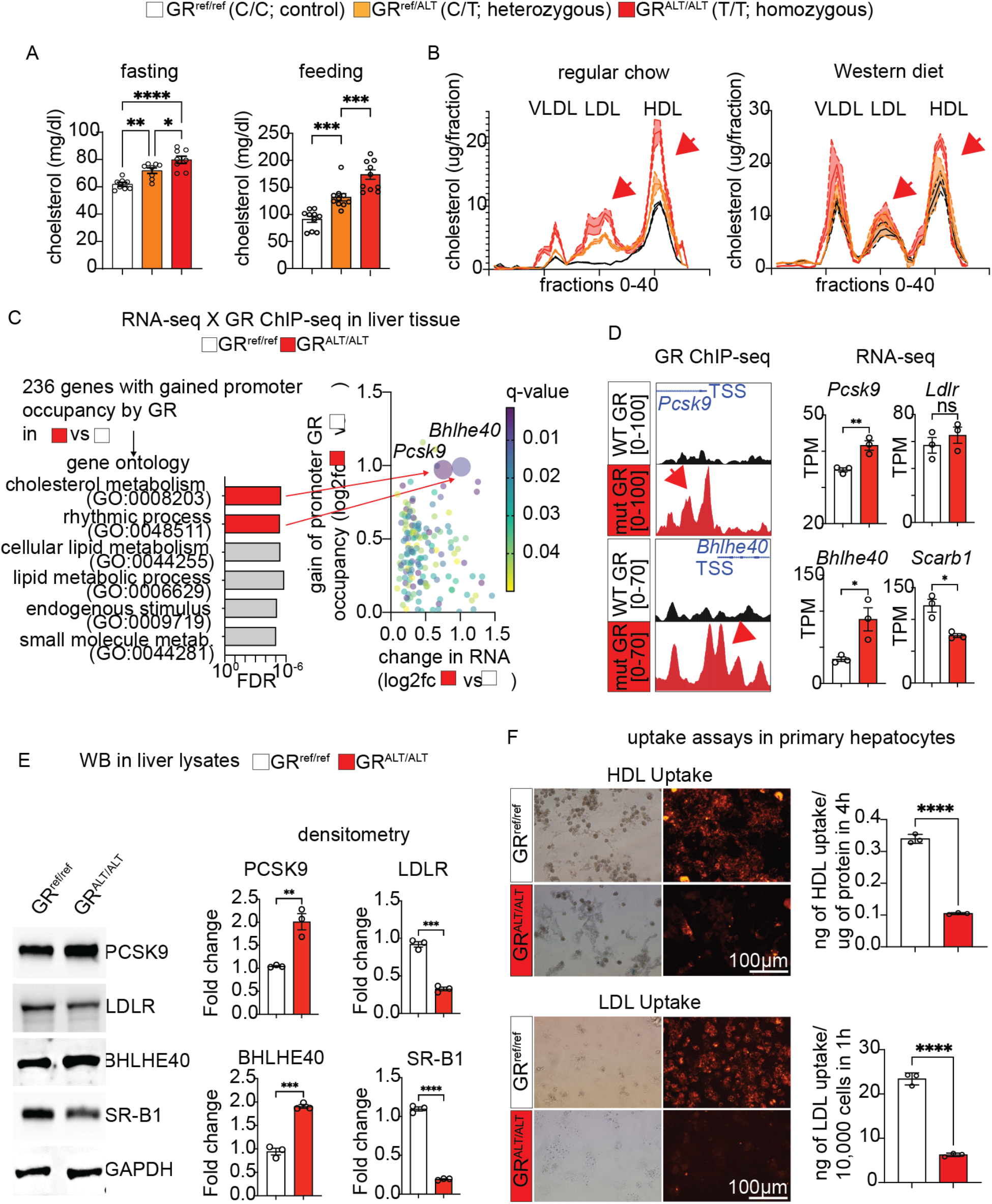
The rs6190 SNP is sufficient to increase cholesterol and skew the liver GR to a gene program repressing liver cholesterol uptake in mice. **(A)** Zygosity-dependent increases in cholesterol in both fed and fasted states in littermates control vs SNP-carrier mice. **(B)** Analogous trends with regular and Western diets, as assayed through FPLC distribution of cholesterol across lipoprotein fractions (arrows highlight increases in LDL- and HDL-cholesterol). **(C)** RNA-seq and ChIP-seq overlay in liver tissue identifies *Pcks9* and *Bhlhe40* as putative transactivation targets of the mutant GR. **(D-E)** ChIP-seq and RNA-seq, as well as validation WB values for PCSK9, BHLHE40 and their putative targets LDLR and SR-B1. **(F)** Uptake of LDL and HDL particles (traced by red fluorescence) is lower in GR^ALT/ALT^ than GR^ref/ref^ primary hepatocytes. N=3-10♀/group, 3-6mo; A: 1w ANOVA + Sidak; D-F: Welch’s t-test; *, P<0.05; **, P<0.01; ***, P<0.001; ****, P<0.0001.

We then focused our mechanistic analyses on GR^ref/ref^ vs GR^ALT/ALT^ liver comparisons, considering the primary role of this organ in cholesterol regulation (24). In line with GTEX (25) predictions on rs6190 as non-significant eQTL for overall *NR3C1* expression levels in liver, overall GR protein levels in liver or primary hepatocytes did not change between GR^ref/ref^ vs GR^ALT/ALT^ mice **(Suppl. Fig. 2D)**. We also checked in other metabolic tissues of primary relevance for lipidemia and cardiovascular health, i.e. adipose tissue and kidney, and found no SNP effects on overall GR protein levels **(Suppl. Fig. 2D)**. As the N-terminus is important for co-factor binding and therefore epigenetic activity of the GR (26), we tested whether the mutant GR had significant changes in interactome by performing GR immunoprecipitation followed by mass-spec (IP-MS) in GR^ref/ref^ vs GR^ALT/ALT^ liver tissues. IP-MS in liver identified Hsp90 among top hits for decreased interaction with the mutant GR compared to the WT GR. CoIP in tissue extracts confirmed decreased GR-Hsp90 interaction in not only liver, but also other metabolically active tissues like adipose and kidney **(Suppl. Fig. 3A)**. Hsp90 is a cytoplasmic docker for the GR and we therefore asked whether the decreased interaction of the mutant GR with Hsp90 corresponded to increased levels of GR nuclear translocation. We tested this in vivo in GR^ref/ref^ vs GR^ALT/ALT^ mice at 30 minutes after a single i.p. 1mg/kg dexamethasone (GR activator) or vehicle pulse, comparing nuclear and cytoplasmic fractions from liver, adipose and kidney tissues. The mutant GR showed increased nuclear/cytoplasmic signal enrichment compared to the WT GR in both control and GR-activated states **(Suppl. Fig. 3B)**, supporting an increased propensity to nuclear translocation by the mutant GR. The changes in liver tissue were recapitulated in primary GR^ref/ref^ vs GR^ALT/ALT^ hepatocytes in vitro for both Hsp90 interaction and nuclear translocation at 30min after 1 µM dexamethasone or vehicle **(Suppl. Fig. 3C)**. Also, in primary hepatocytes from GR^ref/ref^ vs GR^ref/ALT^ vs GR^ALT/ALT^ mice, the mutant GR showed an increased epigenetic activity both at baseline and after dexamethasone stimulation, assayed through a luciferase reporter of direct GR-driven transactivation (**Suppl. Fig. 3C**). Because these data pointed at a SNP-dependent change in GR epigenetic activity, we conducted RNA-sequencing and GR ChIP-sequencing in liver to identify potential differential targets of GR transactivation based on the GR SNP genotype. The liver GR ChIP-seq was validated by enrichment for the canonical GRE motif in unbiased motif analysis (**Suppl. Figure 3D**). Compared to the control GR, the increased epigenomic activity of the mutant GR was evidenced by increased GR signal on GRE sites genome-wide and on the *Fkbp5* promoter, a canonical marker for GR activity (27, 28) (**Suppl. Fig. 3E-F**). No statistical differences were noted in overall peak number or genomic peak distribution, which clustered preferentially in proximal promoter regions for both genotypes (**Suppl. Fig. 3G-H**), suggesting that the mutant GR showed an increased activity on “normal” GR sites, rather than a profound redistribution/acquisition of GR peaks. Liver RNA-seq revealed 368 genes with differential expression by the mutant GR (**Suppl. Fig. 3I**). The overlay of both datasets unveiled 236 genes exhibiting both differential expression and a gain of mutant GR signal on their promoters (**Figure 2C**). Gene ontology (GO) analysis revealed a significant enrichment for cholesterol metabolism. Notably, within this pathway, proprotein convertase subtilisin/kexin type 9 (*Pcsk9*) was the highest hit. The increased transactivation of *Pcsk9* in liver by the mutant GR was validated at mRNA and protein levels (**Figure 2D-E**). Besides indirect and direct inhibition of VLDL-cholesterol clearance (29, 30), PCSK9 plays a pivotal role in increasing circulating LDL cholesterol by promoting the degradation of the main LDL-cholesterol receptor, LDLR, at the protein level (31, 32). Accordingly, the gain in PCSK9 levels correlated with a reduction in protein but not mRNA levels of LDLR in GR^ALT/ALT^ compared to GR^ref/ref^ liver tissues (**Figure 2D-E**). Additionally, within the “rhythmic process” pathway from the ChIP-seq/RNA-seq overlay, the top hit for mutant GR transactivation was *Bhlhe40* (**Figure 2C**), a transcriptional repressor involved in many processes including circadian clock homeostasis (33, 34). Using an ENCODE-mining platform for transcription factor target prediction (35), we screened for putative *Bhlhe40* targets in the promoters of down-regulated genes in mutant versus WT livers. This analysis revealed Scavenger Receptor Class B Type I (SR-B1), encoded by *Scarb1*, as a unique hypothetical target of BHLHE40 from our RNA-seq datasets. SR-B1 is the main receptor for reverse HDL-cholesterol transport in the liver (36). Consistent with our prediction, *Bhlhe40* upregulation correlated with SR-B1 downregulation at both mRNA and protein levels in GR^ALT/ALT^ compared to GR^ref/ref^ liver tissues (**Figure 2D-E**). Additionally, to confirm the in-silico prediction of SR-B1 transcriptional repression by BHLHE40, we compared *Scarb1* expression and SR-B1 protein levels in liver tissues from *Bhlhe40*^*null/null*^ (37) *(Bhlhe40-KO*) vs their wild-type littermate controls (*Bhlhe40-WT*). As hypothesized, SR-B1 was upregulated in the *Bhlhe40-KO* livers compared to WT controls (**Suppl. Fig. 2L**). We then asked the extent to which the mutant GR effect on LDLR and SR-B1 downregulation was biologically significant on hepatocyte biology. We probed fluorescently-labeled LDL and HDL uptake assays in primary hepatocytes to assess this propensity in the absence of body-wide confounders. In line with the LDLR and SR-B1 changes, the GR^ALT/ALT^ hepatocytes showed decreased LDL and HDL uptake *in vitro* compared to GR^ref/ref^ control hepatocytes (**Figure 2H**). Collectively, our findings support a mechanism for the rs6190 SNP effect on cholesterol through which the SNP skews the hepatic GR epigenetic activity and promotes transactivation of *Pcsk9* and *Bhlhe40*, which in turn decreases LDL and HDL cholesterol uptake in liver by repressing LDLR and SR-B1 levels respectively.

### CRISPR-engineered hiPSC-derived hepatocytes confirm the mouse-to-human relevance for the SNP mechanism

In tandem with our murine mouse studies, we questioned whether the molecular SNP mechanism identified was translatable to human hepatocytes. We, therefore, generated SNP heterozygous and homozygous lines from the same parental GR^ref/ref^ hiPSC line through a CRISPR-knockin system. Individual founding clones of isogenic GR^ref/ref^ (control), GR^ref/ALT^ (het), and GR^ALT/ALT^ (homo) hiPSCs were verified through Sanger sequencing and quality-controlled for pluripotency marker expression (**Figure 3A; Suppl. Fig. 4A**). Despite no differences in pluripotency markers, the SNP significantly skewed the GR to a higher rate of glucocorticoid-driven GR translocation in hiPSCs, as shown by serial imaging after a dexamethasone pulse (**Suppl. Fig. 4B**) and consistent with our previous findings with the mutant GR in murine hepatocytes luciferase assay and liver ChIP-seq.

**Figure 3.**
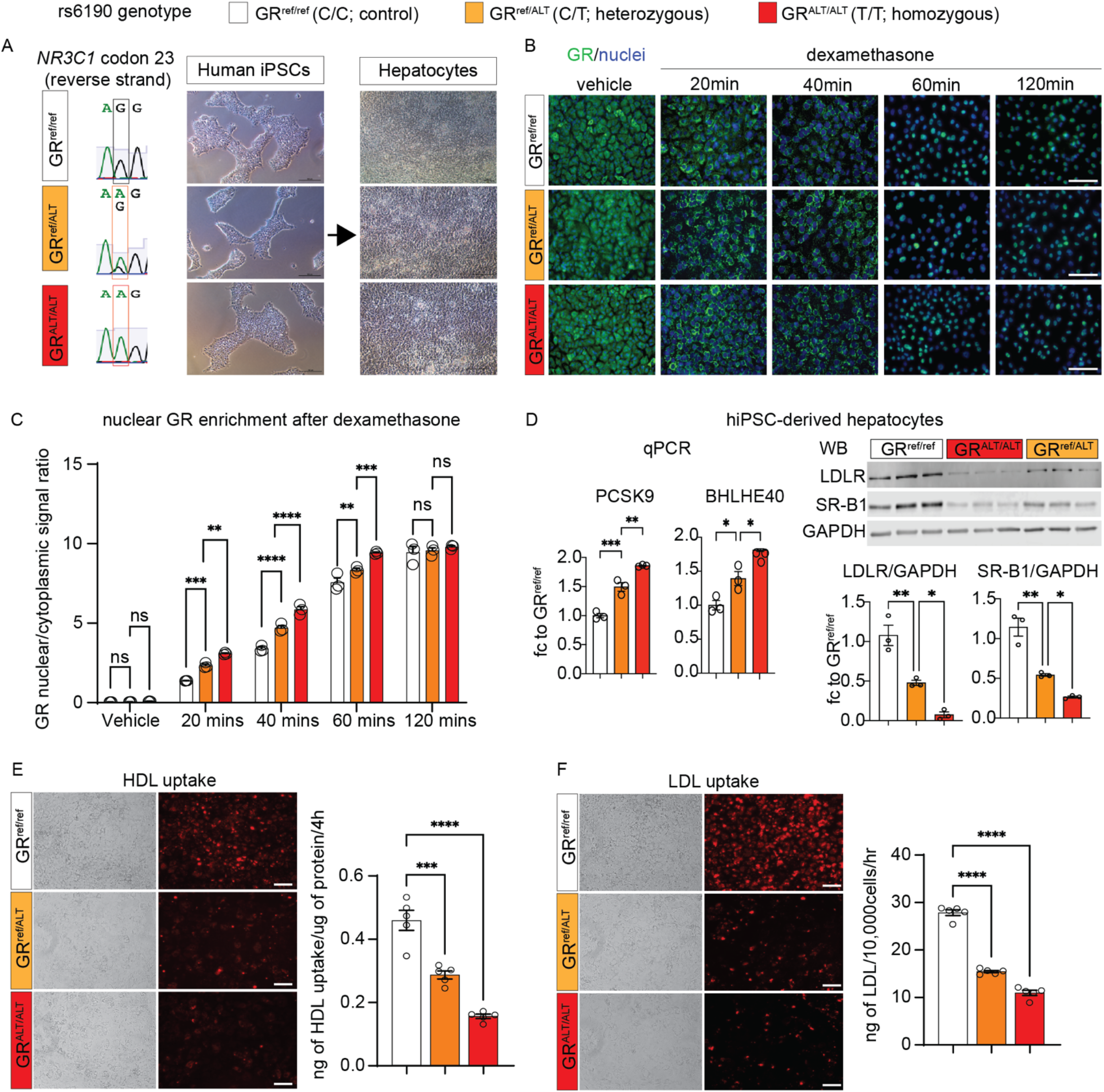
The SNP molecular effects are replicated in hiPSC-derived hepatocytes. **(A)** Sanger sequencing of SNP genotype and brightfield representative images for isogenic hiPSCs and derived hepatocytes with no, one or two rs6190 SNP alleles. **(B-C)** Rate of GR nuclear signal enrichment in hiPSC-hepatocytes increased between 20-60min after dexamethasone addition according to SNP zygosity. **(D)** Zygosity-dependent effects on *PCSK9* and *BHLHE40* upregulation at the hepatocyte level, as well as on protein level downregulation for LDLR and SR-B1. **(E-F)** SNP zygosity replicated the effects on HDL and LDL fluorescent particle uptake in hiPSC-hepatocytes. Scale bars, 100 μm. Each dot represents an independent differentiation replicate; N=3-6/group. B: 2w ANOVA + Sidak; C-E: 1w ANOVA + Sidak. *, P<0.05; **, P<0.01; ***, P<0.001; ****, P<0.0001.

To investigate whether the SNP-mediated molecular mechanism was conserved in human hepatocytes, we subjected the isogenic lines of hiPSCs to a 23-day differentiation protocol to generate mature hepatocyte-like cells (HLCs) (38). Given the well-established role of GR as a regulator of hepatocyte differentiation and maturation (39-41), we sought to investigate whether the presence of the GR SNP influenced the differentiation process. To address this, we examined the expression profiles of differentiation markers at multiple time points during the differentiation process: *NANOG* and *OCT4* at the pluripotent stage (42); *SOX17* and *FOXA2* at the definitive endoderm stage (43); *AFP* and *HNF1A* at the immature hepatocyte stage (44); *ALB* and *CY18*, morphology, and albumin secretion at the mature hepatocyte stage (45). We did not detect any SNP-driven significant alterations in the *in vitro* maturation process of hiPSC-derived hepatocytes (**Suppl. Fig. 4C-D**). However, the hiPSC-derived hepatocytes reproduced the zygosity-dependent increase in GR nuclear translocation (**Figure 3B-C**) and the SNP-mediated effects on *PCSK9* and *BHLHE40* transactivation, as well as post-translational repression of LDLR and SR-B1 (**Figure 3D**). Furthermore, the GR^ALT/ALT^ hiPSC-derived hepatocytes displayed decreased uptake of HDL and LDL-cholesterol compared to GR^ref/ref^ control cells (**Figure 3E-F**). Taken together, our hiPSC-derived hepatocyte data confirm that the molecular SNP mechanism is conserved in human cells and appears autonomous to hepatocytes in the absence of *in vivo* body-wide physiology.

### rs6190 GR SNP promotes atherosclerosis in vivo

Despite our results so far linking the mutant GR to cholesterol regulation, the extent to which the overall SNP-enabled program significantly impacts atherosclerosis *in vivo* remains unknown. To evaluate the extent to which the rs6190 SNP contributes to atherogenic risk *in vivo* in conditions of genetic homogeneity, we crossed our mutant SNP mice with the atherogenic background characterized by homozygous expression of the human *APOE*2* variant (46, 47). The h*APOE*2/*2* mice are well-established transgenic mice known for their susceptibility to atherosclerosis while maintaining cholesterol distribution across all three major lipoprotein compartments (47, 48), unlike other atherogenic backgrounds like *ApoE-KO*. We also excluded the *Ldlr-KO* background as a direct genetic confounder of our LDLR-involving hypothesis.

For these analyses, we focused on GR^ALT/ALT^ vs GR^ref/ref^ female mice. On the h*APOE*2/*2* background and regular chow diet, GR^ALT/ALT^ mice exhibited elevated levels of VLDL-, LDL- and HDL-cholesterol in the FPLC curves compared to control littermates, and this was reinforced even more after a 16-week-long Western diet exposure (**Figure 4A**). We focused on mice exposed to Western diet for atherosclerotic plaque analyses. Compared to GR^ref/ref^, GR^ALT/ALT^ mice exhibited a significant increase in atherosclerotic plaque incidence as quantitated through overall plaque/total aorta area ratio in en face whole aorta staining and imaging (**Figure 4B, left**). Furthermore, histological analysis of the aortic root cross-sections and Oil Red O staining revealed a significant increase in atherosclerotic lesion size (plaque/lumen ratio) and lipid accumulation in GR^ALT/ALT^ versus GR^ref/ref^ mice (**Figure 4B, right**). Finally, considering our hypothesis of *Pcsk9* and *Bhlhe40* as mechanistic mediators of the SNP effect, we tested the effect of *in vivo* knock-down of these genes on the SNP-mediated effect on cholesterol and atherosclerosis through AAV8 vectors. For *Pcsk9* knockdown we used a previously reported AAV vector (49) and confirmed its max knockdown effect in liver in vivo in Apo*2/*2 mice on Western Diet with a 10^13vg/mouse dose **(Suppl. Fig. 5A)**. For *Bhlhe40*, we combined two AAVs with different shRNAs under the U6 promoter, as they showed synergistic effect on *Bhlhe40* knockdown in Apo*2/*2 livers **(Suppl. Fig. 5B)**. At 2 months of age, GR^ALT/ALT^ vs GR^ref/ref^ female mice on the ApoE*2/*2 background were injected retro-orbitally (r.o.) with 3×10^13^ vg/mouse AAV-scramble or 10^12^ vg/mouse/vector AAV-antiPcsk9 (1 vector) + AAV-antiBhlhe40 (2 vectors) immediately before starting the 16-week-long Western Diet exposure. At endpoint, we validated knockdown of PCSK9 and BHLH40, and consistent increases in LDLR and SR-B1 **(Fig. 4C)**, and then focused on FPLC cholesterol curves and atherosclerotic plaques as read-outs. Compared to scramble, the knockdown vectors reduced the cholesterol levels across lipoprotein fractions in GR^ALT/ALT^ mice to GR^ref/ref^-like levels **(Fig. 4D)**, and blunted the SNP-mediated effect on plaque incidence (**Fig. 4E**) and severity **(Fig. 4F)**. We also noted that the knockdown vectors reduced VLDL-cholesterol and plaque incidence but not histological plaque severity in GR^ref/ref^ mice compared to scramble. Taken together, our findings demonstrate that the rs6190 SNP promotes hypercholesterolemia and atherosclerosis *in vivo* through upregulation of *Pcsk9* and *Bhlhe40* in liver.

**Figure 4.**
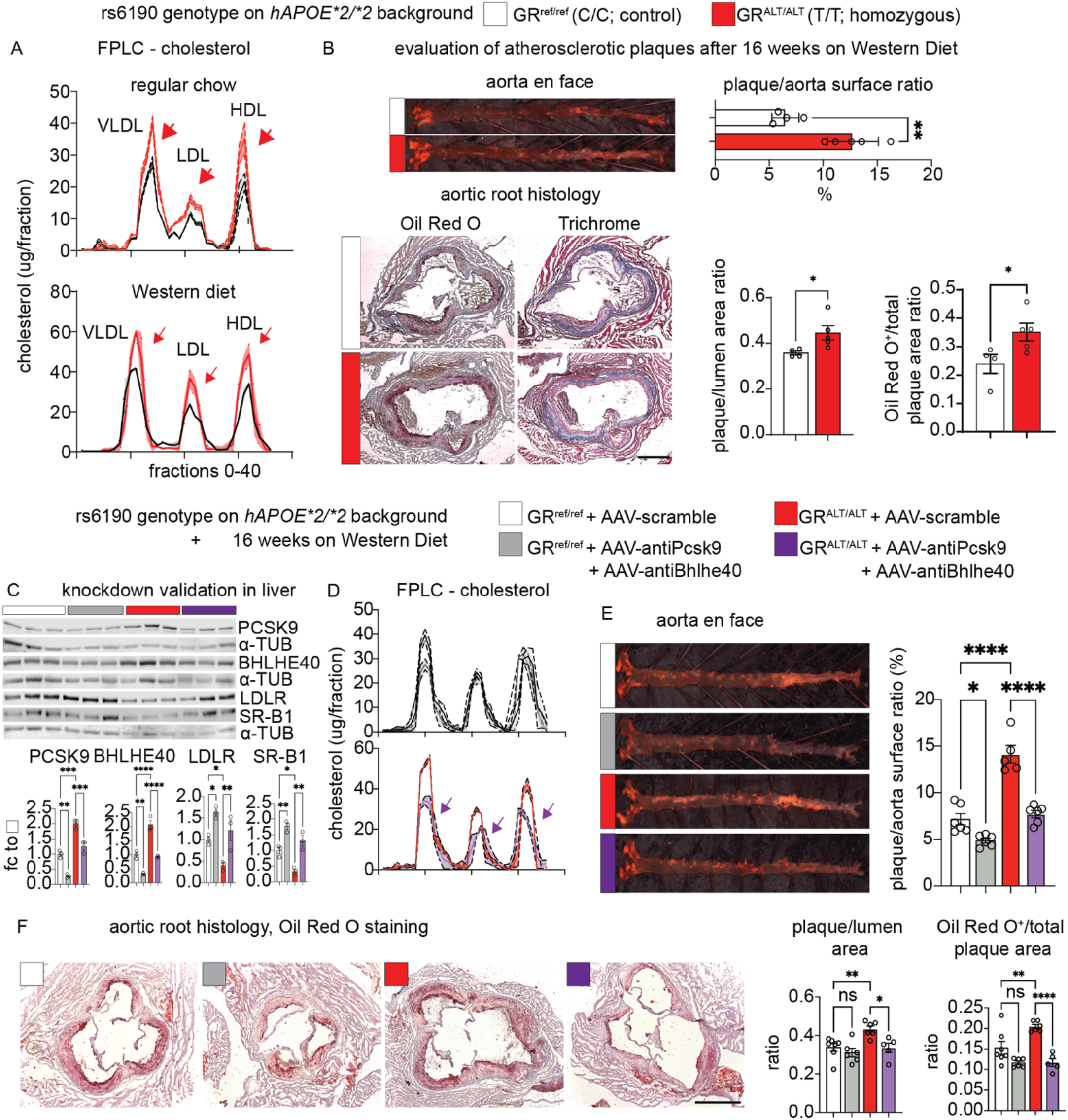
The SNP promotes atherosclerosis in vivo. **(A)** FPLC curves show the additive effect of SNP genotype on the hAPOE*2/*2-driven hypercholesterolemia across lipoprotein fractions in both normal and Western diets (arrows). **(B)** Compared to GR^ref/ref^ mice, GR^ALT/ALT^ mice on the *hAPOE*2/*2* background showed higher incidence (as quantitated from en face analyses) and severity (as quantitated through Oil Red O staining in aortic root sections) of atherosclerotic plaques. **(C)** WB validation of target knockdown in liver. **(D-F)** AAV-mediated knockdown of *Pcsk9* and *Bhlhe40* in adult mice blunted the SNP effect on VLDL-, LDL- and HDL-cholesterol (FPLC), plaque incidence in en face aorta assays, and histological severity of aortic root plaques. Scale bars, 500 μm. N=4-7♀/group, 6mo; B: Welch’s t-test; E-F: 2w ANOVA + Sidak; *, P<0.05; **, P<0.01; ***, P<0.001; ****, P<0.0001.

### Determinants of sexual dimorphism in the SNP effect

Our analyses in human datasets and mice pointed at sexual dimorphism in the rs6190 SNP effect on cholesterol regulation: significant in women and female mice, while not significant in men and male mice. We sought to test the hormonal determinants of this dimorphism and tested two complementary hypotheses in our mutant and control mice: 1) lower corticosterone-driven activation of the mutant GR program in males vs females, and 2) protective effect of testosterone against the “default” SNP effect on cholesterol. For the first question, we measured peak serum corticosterone at ZT16 through ELISA and found modestly but significantly lower corticosterone in males compared to females regardless of SNP genotypes **(Fig. 5A)**, in line with prior sex comparisons for murine corticosterone (50). We reasoned that the lower corticosterone in males could reduce the magnitude of the mutant GR-driven transactivation program in liver and therefore measured the protein levels of the identified cascade (increased PCSK9 and BHLHE40, decreased LDLR and SR-B1) in liver tissues across sexes and genotypes in head-to-head comparisons. The magnitude of the SNP effect on each protein change (GR^ALT/ALT^ vs GR^ref/ref^) was indeed smaller in males than females **(Fig. 5B)**. Albeit modest, the molecular SNP effect was still significant in male livers, consistent with the intrinsic changes induced by the SNP in the GR transactivation program *per se* **(Fig. 5B)**. For the second question, we challenged mutant and control littermate males on either normal (WT background + regular chow) or atherogenic conditions (*hAPOE*2/*2* background + Western diet) with bilateral orchiectomy (ORX). As parallel experimental counterpart and to gain insight into the extent to which the SNP effect is potentially additive to menopause in women, we also challenged female littermates to bilateral ovariectomy (OVX). Sham surgeries were used as control in both sexes. We confirmed the expected testosterone depletion with ORX and estradiol depletion with OVX through serum ELISA at 2-weeks post-surgery in WT mice on regular chow **(Fig. 5C)**. We profiled cholesterol across lipoprotein fractions through FPLC, as this was a significant SNP effect in female mice in both normal and atherogenic conditions. In male mice at 4-months after surgery, the SNP effect was not present in sham-operated mice (no difference in GR^ALT/ALT^ vs GR^ref/ref^), whereas it was present and significant in ORX mice (increased cholesterol across fractions in GR^ALT/ALT^ vs GR^ref/ref^; pink arrows) in both normal and atherogenic conditions **(Fig. 5D)**. ORX per se increased overall cholesterol levels, in line with prior studies in *Ldlr-KO* mice (51). In female mice at 4-months after surgery, the SNP effect was recapitulated in sham-operated mice and additive to the OVX-driven increase in cholesterol across fractions (red and pink arrows; **Fig 5D**). Also in this case, ovariectomy increased cholesterol levels per se compared to sham, in line with prior reports (52). We also analyzed plaque incidence as Oil Red O-based plaque/aorta surface ratio in en face aortas in atherogenic conditions. In males the SNP effect was present only after ORX (pink arrows), while in females the SNP effect was recapitulated in sham and additive to OVX (red and pink arrows; **Fig 5E**). Finally, we counter-tested the overall endocrinological logic behind our sexual dimorphism hypothesis in our hiPSC-derived hepatocytes. Our GR^ALT/ALT^ and GR^ref/ref^ hiPSCs are all derived from the parental hiPSC line 72.3, which is a male cell line generated by the CCHMC Pluripotent stem cell facility from a male donor’s foreskin (CCHMCi001-A cell line on Cellosaurus) with a normal human male karyotype shown in the original publication describing this line (53). We also reconfirmed the male cellular sex in hiPSC-hepatocytes by verifying higher expression of androgen over estrogen receptor by qPCR (*Ar, Esr1* genes; **Fig. 5F**). Considering these cells are grown and differentiated as naïve to sex hormones in vitro, we asked whether testosterone could oppose the SNP effect in vitro. We focused on HDL and LDL uptake, both of which were significantly impacted by the SNP effect in our prior findings in hiPSC-hepatocytes at baseline, i.e. in the absence of specific challenges. We tested HDL and LDL uptake at 24-hours after exposure to 100nM CL-4AS1, a steroidal androgen receptor agonist (54), or 100nM β-estradiol, or vehicle. The SNP-driven loss of HDL and LDL uptake was recapitulated in the vehicle-treated hepatocytes (lower in GR^ALT/ALT^ and GR^ref/ref^), yet CL-4AS1 increased uptake of both particle types in both genotypes compared to vehicle, confirming the notion that testosterone-driven effects are independent from the genotype-driven effects in hepatocytes at the molecular level. We did not find significant changes with β-estradiol in these acute settings. Taken together, these results indicate that lower corticosterone and testosterone protect males from the SNP-driven elevation of cholesterol and atherosclerosis, while that SNP effect is additive to estrogen loss in females.

**Figure 5.**
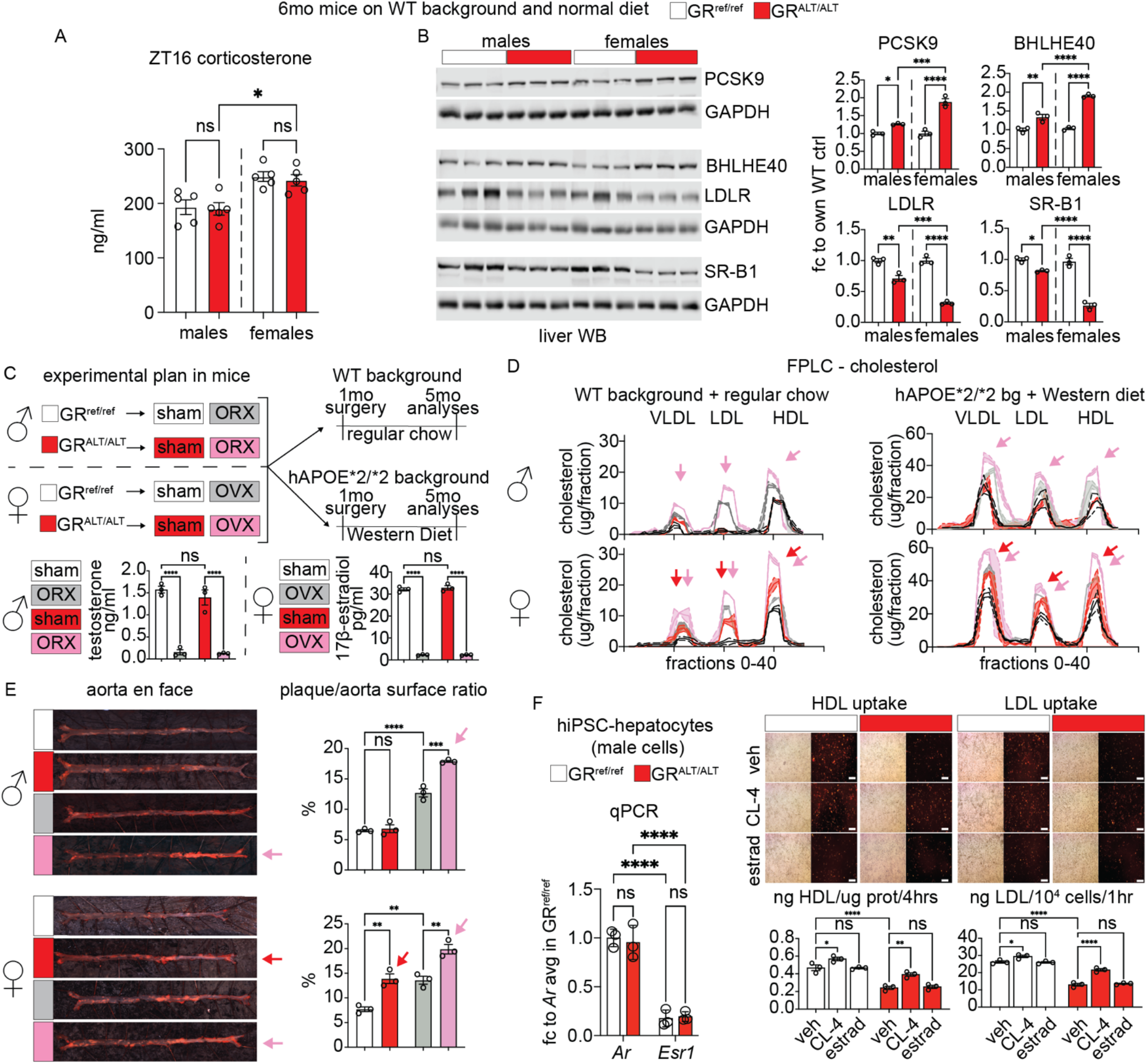
Hormonal determinants of sexual dimorphism in the SNP effect. **(A)** Serum corticosterone at its typical peak circadian time in mice ZT16 was assessed by ELISA and was higher in females than males of both control and mutant genotypes. **(B)** WB analyses in liver tissue showed that the molecular SNP effect on the identified protein cascade (increased PCSK9 and BHLHE40, decreased LDLR and SR-B1) is present in males but the effect magnitude is significantly higher in females. **(C)** Experimental plan for gonadectomy experiment with ELISA-based validation of serum testosterone or estradiol depletion in WT mice on regular chow at 2-weeks post-surgery. **(D)** In male mice, orchiectomy unmasked the SNP effect on increased cholesterol across lipoprotein fractions in either normal (WT background + regular chow) or pro-atherogenic (hAPOE*2/*2 background + Western diet) conditions (pink arrows). In females, the SNP effect was recapitulated in sham mice (red arrows) and maintained after ovariectomy (red arrows). Y-axes scales are different between normal and atherogenic conditions to adapt to the different cholesterol levels. **(E)** In atherogenic conditions, orchiectomy unmasked the SNP effect on male plaque/aorta surface ratio (pink arrows), while in females the SNP effect was additive to the ovariectomy effect on the aorta plaque profile (red and pink arrows). **(F)** Male hiPSC-hepatocytes showed higher mRNA expression of the AR than the ER per qPCR. The hepatocyte-autonomous protective effect of testosterone on HDL or LDL uptake was modest but significant in vitro in both control and mutant genotypes. We did not detect significant changes induced by estradiol. Scale bars, 100 μm. N=3-5/sex/group; 2w ANOVA + Sidak; *, P<0.05; **, P<0.01; ***, P<0.001; ****, P<0.0001. bg, background; CL-4, CL-4AS1, AR agonist; estrad, β-estradiol.

## Discussion

The glucocorticoid receptor (GR) is well-known for its involvement in orchestrating large gene programs and modulating hepatic lipid and glucose metabolism. However, the precise mechanisms by which hepatic GR governs cholesterol regulation remains elusive. Despite the well-established association between chronic glucocorticoid exposure and hypercholesterolemia with concomitant metabolic stress (55), a direct link between GR and atherosclerosis remains unclear. In this study, we leveraged a naturally occurring human mutation, the rs6190 SNP, to unveil a direct GR-mediated program governing hepatic cholesterol regulation and its consequential implication for atherogenic risk. We focused here on the hepatic transactivation targets of the mutant GR based on ChIP-seq-RNA-seq overlay, and consequently validated *Pcsk9* and *Bhlhe40* as mediators of the SNP effect on LDLR and SR-B1 levels in liver, as well as on overall cholesterol levels and atherosclerosis in the *hAPOE*2/*2* background. We recognize that our study did not address potential mutant GR effects on apolipoproteins (e.g. ApoE itself) or macrophages, both critical determinants of atherosclerosis and in turn regulated by glucocorticoids and/or GR (13, 14). While beyond the focus of the present study, these are compelling questions to address to expand significance of our findings for overall hypercholesterolemia and atherosclerosis risk in SNP carriers.

Our mixed-model regressions in the UK Biobank and the All Of Us datasets have unveiled an unexpected association between the rs6190 SNP and modest but significant elevations of total, LDL-, and HDL-cholesterol in women. Importantly, the impact of the rs6190 genetic variant demonstrated an additive effect based on SNP zygosity, i.e. according to the number of “risk” alleles. Additionally in the UK Biobank, the rs6190 SNP correlated with increased odds ratio for hypercholesterolemia and cardiovascular-related mortality. It was compelling to find analogous correlations in two cohorts that are quite different with regards to genetic ancestry composition. In the All Of Us cohort, the highest minor allele frequency for the SNP was in individuals with European ancestry and closely matched the minor allele frequency of the UK Biobank, where the “white British ancestry” indeed accounts for almost 90% of the cohort (56). Beyond SNP correlations in human datasets, we sought to gain the mechanistic insight in mice and hiPSCs of the extent to which the rs6190 SNP is sufficient to regulate cholesterol. Our findings in murine liver and hiPSC-derived hepatocytes show the SNP is indeed sufficient to elevate cholesterol and promote atherosclerosis through a specific change in the GR activity. In principle, this is a novel mechanism of SNP action that is independent from the genomic context, and future studies will be needed to articulate the genetic modifiers that potentiate or contrast this mechanism across ancestries in the human population. Variants in the *NR3C1* gene are generally not significant enough to associate with cholesterol changes in GWAS studies with a typical p-value threshold of 5×10-8. Indeed, variants like rs6190 are low-frequency or rare and with modest effects on complex metabolic traits. Our study combined candidate variant analyses with rigorous mechanistic dissection in mice and hiPSCs to “unmask” the rs6190 role in GR biology and cholesterol regulation. It will be intriguing in the future to use analogous study settings to delve into the non-coding variants we found associated with total cholesterol in the All of Us, and to dissect whether they influence the rs6190 SNP effect as genetic modifiers or in haplotypes.

Given the well-established role of GR as a potent transcription factor, we examined the potential alterations in the epigenetic activity of GR induced by the rs6190 mutation. At the molecular level, our findings revealed that the mutant GR exhibited increased epigenetic activity and nuclear translocation, leading to the differential expression of 236 genes, including key regulators of cholesterol metabolism. Notably, the mutant GR upregulated *Pcsk9*, a key regulator of VLDLR and LDLR degradation, and *Bhlhe40*, a circadian transcriptional repressor that is implicated in SR-B1 control. At present, additional experiments are required to ascertain the extent to which the increase in cholesterol is independent of general changes in lipidemia. However, we emphasize that our regression analyses in women from the UK Biobank dataset took into account triacylglycerols as co-variate and still found a significant zygosity-dependent effect on total and LDL-cholesterol. We also wish to note that the knockdown experiment with Pcsk9 and Bhlhe40 in SNP-bearing atherogenic mice blunted but did not completely remove the SNP effect on hypercholesterolemia and plaques. This suggests additional pathways elicited by rs6190 in atherosclerosis risk regulation that future studies will have to dissect.

To confirm the conservation of the SNP-mediated mechanism, we utilized isogenic hiPSC-derived hepatocytes carrying the rs6190 SNP. These hiPSC-derived hepatocytes exhibited increased expression of *PCSK9* and *BHLHE40*, consistent with murine model findings. Moreover, these hepatocytes demonstrated reduced uptake of HDL and LDL cholesterol, providing direct evidence that the SNP influences cholesterol regulation and this mechanism is conserved in human cells. Although the rs6190 SNP is described in ClinVar as associated with “glucocorticoid resistance,” our analyses in hiPSCs and hiPSC-derived hepatocytes revealed that the mutant GR is more susceptible to glucocorticoid-induced activation than the reference GR isoform. This observation suggests that the SNP may confer increased “glucocorticoid sensitivity” in addition to its effects on cholesterol regulation. The evidence in support of “glucocorticoid resistance” is mostly limited to one study, where targeted limited analyses found that rs6190 decreased dexamethasone-driven activation of *GILZa* in immune cells (57). However, several subsequent studies have failed to find correlation between rs6190 and reduced sensitivity to glucocorticoids, including the seminal study that first discovered the rs6190 polymorphism (22, 27, 55, 58-61). Further *in vitro* experiments are warranted to investigate the extent to which the mutant GR activates newly identified glucocorticoid response elements (GREs) dependently or independently from other key nuclear factors for cholesterol regulation.

Finally, our results with corticosterone and gonadectomies in vivo indicate that the sexual dimorphism in the rs6190 effect on cholesterol and atherosclerosis elevation (higher in women and female mice than in men and male mice) results from the interaction between a sex-independent molecular program and a sex-specific endocrinological effect. The SNP effect on the mutant GR transactivation program is sex-independent, as shown by target liver protein changes in both sexes in mice. However, the “conversion” of this program into actual changes in cholesterol and atherosclerosis depends on the superimposed endocrinological signals of corticosterone and sex hormones. Corticosterone is typically higher in female mice compared to male mice without specific stress challenges (50), and this could find a parallel in humans, as shown by an observational study with salivary cortisol in 1671 people that reported higher morning-cortisol and delta-cortisol (morning - evening cortisol) levels for women than men in the absence of known specific stressors (62). In our mice, the higher corticosterone in females was additive to the SNP-based change in liver GR transactivation program compared to males. This finding opens the idea of corticosterone as potential new “dimorphic pressure” to consider in the case of sex-different GR processes beyond cholesterol. With regards to cholesterol, our findings with gonadectomies are consistent with the reported protective effects of testosterone (51) and estrogens (52) in atherosclerosis. In the rs6190 case, testosterone appeared to work “against” the SNP program and indeed testosterone removal through orchiectomy “unmasked” the SNP effect on cholesterol and atherosclerosis in male mice in vivo. Accordingly, a testosterone-mimic (CL-4AS1) partially rescued the HDL-LDL uptake defect in male hiPSC-hepatocytes. We used male hiPSC-hepatocytes as all our SNP-mutant lines were CRISPR-derived from the same parental male line. Nonetheless, as hiPSC-hepatocytes are grown and differentiated outside the typical physiological sexual development of a body, the cells recapitulated the SNP-driven GR transactivation cascade at the molecular level in vitro. Future comparisons with male and female hiPSC lines – ideally from different genetic backgrounds – are required to resolve the extent to which chromosomal sex impacts rs6190 effects in pluripotent and differentiated states. Our findings with corticosterone and testosterone are consistent with the seminal finding of a cholesterol signal for rs6190 in women but not men of both the UK Biobank and the All of Us cohorts. They are also consistent with a previous study that found a significant association between rs6190 and sexually dimorphic cholesterol changes in a small cohort with heterozygous SNP carriers (18). We also found that estrogen loss through ovariectomy is additive to the SNP effect on cholesterol and plaque incidence in female mice. This finding suggests that the SNP genotype could interact with menopause as additive atherosclerosis risk in older women, but specific studies are required to districate that interaction from social determinants of health and other complex variables like genetic ancestry, aging-related co-morbidities, hormonal interventions/variations and lifestyle.

### Limitations of this study

Besides specific limitations and considerations reported above for specific results, in this study we have not formally assessed the impact of the single amino acid change (R->K) on the N-terminus structure or the overall conformation of the GR. Moreover, based on the initial cholesterol signal in the human datasets, we focused on liver GR and cholesterol. However, the SNP will expectedly impact the GR in virtually every tissue and additional studies are required to dissect the extent and mechanisms of rs6190-driven changes in the physiological GR action in other tissues. Lastly, while focusing on the upstream genetic, epigenetic and molecular mechanisms related to cholesterol regulation by the GR, we recognize that our study did not address the translational question of whether rs6190 impacts efficacy of the current strategies for cholesterol lowering.

### Conclusions and overall impact

In conclusion, our study leverages the rs6190 SNP as genetic linchpin to advance our understanding of the GR-driven regulation of cholesterol through genetic and epigenetic mechanisms. Our data support early and proactive monitoring for cholesterol in carriers of this non-rare variant, particularly in women.

## METHODS

Methods for circulating cholesterol, hormones, RNA-seq, ChIP-seq, WB, hiPSC maintenance, albumin, hepatocyte isolation, immunostaining, nuclear/cytoplasmic fractions, co-IP are available as Supplementary Information.

### Sex as biological variable

Our seminal screening of UKB NMR metabolomics datasets was performed disaggregated by sex considering the known outsized role of sexual dimorphism on virtually all listed metabolic parameters. Subsequent correlations and zygosity stratifications were also then performed as disaggregated by sex and data from both sexes are reported. In mice, analyses were also performed disaggregated by sex and data from both males and females are presented. For hiPSC-based experiments, only male cells were used as all our control and mutant cell lines were derived from the seminal male hiPSC line CCHMCi001-A (RRID:CVCL_A1BW). Despite this limitation, these cells were able to model in vitro the molecular SNP effect on the GR transactivation cascade.

### Animals and diet

Mice used in this study were maintained in a pathogen-free facility in accordance with the American Veterinary Medical Association (AVMA) and under protocol fully approved by the Institutional Animal Care and Use Committee (IACUC) at Cincinnati Children’s Hospital Medical Center (#2023-0002). Euthanasia of the mice was carried out in accordance with ethical guidelines. Carbon dioxide inhalation was utilized as the initial method for euthanasia, followed by cervical dislocation and removal of the liver tissue.

All animals were maintained in a temperature-controlled environment with a 12h/12h light/dark cycle. For the fasting group, mice were subjected to an 18-hour starvation period. Mutant GR mice were generated using CRISPR/Cas9 genome editing by genocopying the rs6190 SNP in the endogenous *Nr3c1* locus on the C57BL/6J background. This genetic modification was performed by the Transgenic Animal and Genome Editing Core Facility at CCHMC. To ensure genetic background homogeneity and control for potential confounding variables, the colonies were maintained through heterozygous matings. This approach allowed us to compare three distinct groups of mice as littermates: GR^ref/ref^ (control WT), GR^ref/ALT^ (heterozygous SNP carriers), and GR^ALT/ALT^ (homozygous SNP carriers). All animals used in this study were approximately 3-4 months of age at the time of experimentation. As the primary atherogenic model, *hAPOE*2/*2* homozygous mice were originally obtained from the Maeda Laboratory at the University of North Carolina (47) and maintained as breeding colony from Dr. David Hui’s lab at the University of Cincinnati. These mice were crossed with the R24K mutant mice. To induce hypercholesterolemia and atherosclerosis, R24K mice crossed on *hAPOE*2/*2* background were subjected to cholate-free western diet, which contained 21% fat and 0.2% cholesterol for 16 weeks.

At one month of age, both male and female GR^ref/ref^ and GR^ALT/ALT^ mice underwent either gonadectomy or sham operations. Anesthesia was induced and maintained with 2% isoflurane. For ovariectomy (OVX) and orchidectomy (ORX), a midline incision was made to access and remove ovaries or testicles, respectively. Sham-operated animals underwent identical surgical procedures without gonadal removal. Blood samples were collected from tail vein sampling for serum isolation and hormone analyses at two-weeks post-surgery. R24K mice on the WT C57BL/6J background were fed with a normal chow diet, while R24K mice on *hAPOE*2/*2* background were administered a western diet. Sixteen weeks post-surgery, mice were fasted overnight and euthanized at ZT4 to control for circadian and feeding variables.

For systemic AAV experiments, wild-type and homozygous SNP-mutant littermate mice on *hAPOE*2/*2* background were injected retro-orbitally with either 3×10^13^ vg/mouse of AAV8-scramble shRNA or 1×10^13^ vg/mouse for each of the knockdown combination vectors, i.e. one AAV8-anti*Pcsk9* (49) and two AAV8-*Bhlhe40*shRNA vectors (Vector Builder vectors # VB010000-0023jze, VB230421-1310pka, VB230421-1312ydp; Addgene #163025; scramble shRNA sequence: CCTAAGGTTAAGTCGCCCTCG; anti-*Bhlhe40* shRNA sequences: GCGAGGTTACAGTGTTTATAT, GTAGTGGTTTGGGCAAATTTC) while under inhaled isoflurane anesthesia. All AAV8 injections were diluted in sterile PBS. To prepare and isolate AAV virions, we followed the procedures we previously reported (63, 64).

### Lipoprotein analysis

For lipoprotein separation through FPLC, fresh plasma samples were pooled, totaling 250 µl, obtained from at least 5 mice per group. Each group’s pooled plasma underwent FPLC gel filtration, utilizing a tandem arrangement of 2 Superose 6 columns (GE Healthcare). The elution process entailed the collection of fractions in 0.5 ml increments, maintaining a steady flow rate of 0.5 ml/min. This procedure yields a total of fifty-one distinct fractions, each of which is subjected to quantification of total triglyceride and cholesterol levels using the Infinity Triglyceride and Cholesterol kits.

### Atherosclerotic lesion analysis

Mice under anesthesia were subjected to a perfusion procedure using a 10% formalin solution in buffered saline for 5 mins. Following this perfusion, the hearts were carefully dissected to harvest aortic roots. These harvested tissues were subsequently preserved in 10% buffered formalin solution. To assess the distribution of atherosclerosis, en face whole aorta lesion staining was performed with Oil Red O for 30 mins, followed by two 1x PBS washes. Furthermore, the aortic root of the heart was embedded in OCT compound for the preparation of frozen sections. Cross cryosections of the aortic roots, measuring 7µm in thickness and encompassing the aortic valve region, were stained with H&E, Oil Red O and Trichrome staining according to our established protocols. Images were obtained using a ZEISS Axio Imager.A2 microscope and histological analyses performed using the ImageJ software (NIH).

### Human iPSC-derived hepatocyte-like cells (HLCs) differentiation in vitro

When human iPSCs reached a confluency of approximately 95% they were passaged with Accutase™ Cell Dissociation Reagent (Cat #07920, StemCell Technologies) and resuspended as single cells in mTesR™1 medium with 10 µM Y-27632 (Tocris Bioscience). The cells were seeded in six well plates pre-coated with Cultrex diluted in ice-cold DMEM/F12 (Thermo Fisher Scientific). After 24 hours, wash the cells with room temperature DMEM/F12 and switch to RPMI 1640 (Cat #11875093, Thermo Fisher Scientific) with B27 supplement Minus Insulin (Cat #A1895601, Thermo Fisher Scientific), along with 100 ng/ml Activin A (Cat #120-14P, Peprotech) and 3 µM CHIR99021 (Cat #4423, Tocris Bioscience). Following 24-h treatment, CHIR99021 was withdrawn, and the cells were treated with RPMI 1690/B27 Minus Insulin basal medium with 100 ng/ml Activin A for another 48 hours and renewed every day to generate definitive endoderm cells (DE). The differentiated endoderm cells were further treated with RPMI 1640/B27 Minus Insulin along with 10 ng/ml basic fibroblast growth factor (FGF) (Cat 3100-18B, Peprotech) and 20 ng/ml Bone morphogenic factor 4 (BMP4) (Cat #120-05ET, Peprotech). The media was replaced every day for the next 5 days to generate hepatic progenitor (HP) cells. Next, the hepatic progenitors were further differentiated into immature hepatocytes (IMH) by replacing the media with RPMI/B27 Minus Insulin, 20 ng/ml hepatocyte growth factor (HGF) (Cat #100-39, Peprotech), and 0.5% DMSO. The media was replaced every day for the next 5 days. To promote maturation of immature hepatocytes, the media was replaced with HCM™ Hepatocyte Culture Medium Bulletkit™ (Cat # CC-3198, Lonza) except HEGF, 10 ng/ml HGF, 20 ng/ml Oncostatin M (Cat #300-10T, Peprotech), 100 nM Dexamethasone (Cat # D2915, Sigma), and 0.5% DMSO for another 5 days with everyday media change.

For the GR translocation assay and analysis, hiPSCs were exposed to either a vehicle control or 1 µM Dexamethasone for various time intervals (20 mins, 40 mins, 60 mins, and 120 mins). Subsequently, an immunofluorescence assay was performed. To evaluate GR translocation in hiPSC-derived mature HLCs, the maturation medium containing 100 nM Dexamethasone was removed, and the cells were cultured in hepatocyte maintenance (HCM) medium without dexamethasone for 24 hours. The following day, mature HLCs were treated with either vehicle control or 1 µM Dexamethasone for the aforementioned time intervals. Immunofluorescent staining was performed using GR (Cat #sc-393232, 1:200, Santa Cruz) and Alexa Fluor® 488 AffiniPure Donkey Anti-Mouse IgG (H+L) (Cat #102650-156, 1:300, VWR). The analysis of GR translocation was carried out using ImageJ software on 5-6 images per sample acquired from a Nikon Eclipse Ti-U microscope.

### Fluorometric HDL and LDL uptake assay and quantitation

Plate 3-4×10^4^ cells/ well in a 96-well white clear-bottom cell culture plates and culture in media overnight at 37°C incubator. Next day, wash the cells with Assay buffer provided in this appropriate kit. For fluorometric HDL (Cat #ab204717, Abcam) and LDL (Cat #770230-9, Kalen Biomedical) staining and quantitation, we followed the instructions according to the manufacturer; protected samples from light and measured the fluorescence in a microplate reader. For hormone treatment experiments, iPSCs were differentiated into hepatocytes in 96-well plates. At the end of differentiation process, cells were treated with vehicle, 100 nM testosterone, or 100 nM estradiol for 24 hrs. Subsequently, cells were washed with Assay buffer, and HDL and LDL uptake assays were performed as described above. All the procedures were conducted in accordance with the manufacturer’s instructions to ensure assay integrity and reproducibility.

### UK Biobank and All of Us analyses

Our analyses were conducted under the UKB application number 65846 and All of Us workspace number aourw-0fb52975. We constructed a rs6190 genotype-stratified cohort, excluding participants if they withdrew consent. All available values for the tested parameters were collected per genotype group. For UK Biobank, UDI and related parameters: Age: 21001-0.0; BMI: 21001-0.0; Glycemia (mM): 30740-0.0; Triglycerides (mM): 30870-0.0; Total Cholesterol: 23400; ICD10 causes of death, primary 40001, secondary 40002. For initial discovery using the NMR metabolomics datasets, quantitative linear regression and conditional analyses were performed using an additive genetic model adjusting for 10 PCs, sex; and age. In conditional analyses, the 12 established SNP dosage effects were also included as additional covariates. Regression analyses were performed using second generation of PLINK (65). Before analyses, a series of standard QC measures were applied including sample call rates, sample relatedness, and sex inconsistency as well as marker quality (i.e., marker call rate, minor allele frequency (MAF), and Hardy-Weinberg equilibrium (HWE). Analyses were limited to participants with call rates > 98%, SNPs with call rates > 99%, and SNPs with MAF > 1% and HWE p > 0.0001. For independent association confirmation studies, multiple linear regression analysis was carried out using R 4.3.2 (R Core Team, 2023) to explore the association of total cholesterol, clinical LDL, and HDL cholesterol versus separate sex (males/females) and correcting for BMI, glycemia, and triglycerides.

### Immunoprecipitation with Bead-Antibody Conjugation and LC-MS/MS Analysis

Immunoprecipitation of proteins, minimizing contamination from antibody heavy and light chains, was performed using the Pierce Co-IP Kit (Invitrogen #26149). Briefly, 10-75 µg of antibody was conjugated to the AminoLink Plus Coupling Resin using a coupling buffer containing sodium cyanoborohydride as the conjugation reagent. The reaction was carried out in a 1.5 mL Eppendorf tube using a thermomixer at room temperature for 2 hours. Simultaneously, protein extracts were pre-cleared with control agarose resins for 1 hour. Following pre-clearing, the resin was washed with a series of quenching and wash buffers. The eluted pre-cleared protein extracts were then incubated with the antibody-conjugated AminoLink resin overnight at 4°C. The next day, the resin was washed with wash buffer, and the bound proteins were eluted using 50 µL of elution buffer. The eluted protein samples were analyzed via SDS-PAGE, silver staining, and western blot to confirm the presence of antibody-free immunoprecipitated proteins. Once validated, the samples were submitted to the LC-MS/MS protein core at UC. For LC-MS/MS analysis, protein samples were dried using a SpeedVac and resuspended in 35 µL of 1X LB. The samples were loaded 1.5 cm into an Invitrogen 4-12% Bis-Tris gel using MOPS buffer, with molecular weight marker lanes included for reference. Gel sections were excised, reduced with DTT, alkylated with IAA, and digested overnight with trypsin. The resulting peptides were extracted, dried using a SpeedVac, and resuspended in 0.1% formic acid (FA). A total of 500 ng to 2 µg of each sample was analyzed by nano LC-MS/MS using an Orbitrap Eclipse mass spectrometer. The data were searched against a combined database of contaminants and the SwissProt Mus musculus database using Proteome Discoverer version 3.0 with the Sequest HT search algorithm (Thermo Scientific).

### Statistics

Unless differently noted, statistical analyses were performed using Prism software v8.4.1 (GraphPad, La Jolla, CA). The Pearson-D’Agostino normality test was used to assess data distribution normality. When comparing the two groups, a two-tailed Student’s t-test with Welch’s correction (unequal variances) was used. When comparing three groups of data from one variable, one-way ANOVA with Sidak multi-comparison was used. When comparing data groups for more than one related variable, two-way ANOVA was used. For ANOVA and t-test analyses, a P value less than 0.05 was considered significant. When the number of data points was less than 10, data were presented as single values (dot plots, histograms). Tukey distribution bars were used to emphasize data range distribution in analyses pooling larger data points sets per group (typically > 10 data points). Analyses pooling data points over time were presented as line plots connecting medians of box plots showing distribution of all data per time points. Randomization and blinding practices are followed for all experiments. All the data from all animal cohorts and cell clone replicates is reported, whether outlier or not.

### Study approval

Mice were housed in a pathogen-free facility in accordance with the American Veterinary Medical Association (AVMA) and under protocols fully approved by the Institutional Animal Care and Use Committee (IACUC) at Cincinnati Children’s Hospital Medical Center (#2022-0020, #2023-0002). UKB and All of Us analyses were conducted under the UKB application number 65846 and All of Us workspace number aou-rw-0fb52975.

## Supporting information

Supplemental Material bundle

## Data availability

RNA-seq and ChIP-seq datasets reported here are available on GEO as GSE280494 and GSE280572. Individual data for all charts presented here is available in the Supporting Data Values file.

## Author contributions

HBD, AH, GN, ADP, KMF, HL, AR, OA, BNK: Data curation, Formal analysis, Investigation; AJ, DYH: Resources; MQ: Conceptualization, Formal analysis, Funding acquisition, Supervision.

## Acknowledgements

Next-gen sequencing was performed thanks to the Cincinnati Children’s DNA Sequencing and Genotyping Facility (RRID: SCR_022630), with critical assistance by David Fletcher, Keely Icardi, Julia Flynn, and Taliesin Lenhart. hiPSC generation, engineering and initial quality control/selection were performed thanks to the Cincinnati Children’s Pluripotent Stem Cell Facility (RRID: SCR_022634), with critical assistance by Chris Mayhew and Yueh-Chiang Hu. pAAV Alb-AAT KRAB-SadCas9 U6-mPcsk9 was a gift from Tonia Rex (Addgene plasmid # 163025 ; http://n2t.net/addgene:163025; RRID:Addgene_163025).

## References

1. El Khoudary SR, Aggarwal B, Beckie TM, Hodis HN, Johnson AE, Langer RD, et al. Menopause Transition and Cardiovascular Disease Risk: Implications for Timing of Early Prevention: A Scientific Statement From the American Heart Association. Circulation. 2020;142(25):e506–e32.

2. Hodgin JB, and Maeda N. Minireview: Estrogen and Mouse Models of Atherosclerosis. Endocrinology. 2002;143(12):4495–4495.

3. Cupido AJ, Asselbergs FW, Schmidt AF, and Hovingh GK. Low-Density Lipoprotein Cholesterol Attributable Cardiovascular Disease Risk Is Sex Specific. J Am Heart Assoc. 2022;11(12):e024248.

4. de Guia RM, Rose AJ, and Herzig S. Glucocorticoid hormones and energy homeostasis. Horm Mol Biol Clin Investig. 2014;19(2):117–117.

5. Schaaf MJ, and Cidlowski JA. Molecular mechanisms of glucocorticoid action and resistance. J Steroid Biochem Mol Biol. 2002;83(1-5):37–48.

6. Lim HW, Uhlenhaut NH, Rauch A, Weiner J, Hubner S, Hubner N, et al. Genomic redistribution of GR monomers and dimers mediates transcriptional response to exogenous glucocorticoid in vivo. Genome Res. 2015;25(6):836–836.

7. Oakley RH, and Cidlowski JA. The biology of the glucocorticoid receptor: new signaling mechanisms in health and disease. J Allergy Clin Immunol. 2013;132(5):1033–1033.

8. Watts LM, Manchem VP, Leedom TA, Rivard AL, McKay RA, Bao D, et al. Reduction of hepatic and adipose Mssue glucocorticoid receptor expression with antisense oligonucleotides improves hyperglycemia and hyperlipidemia in diabetic rodents without causing systemic glucocorticoid antagonism. Diabetes. 2005;54(6):1846–1846.

9. Petrichenko IE, Daret D, Kolpakova GV, Shakhov YA, and Larrue J. Glucocorticoids stimulate cholesteryl ester formation in human smooth muscle cells. Arterioscler Thromb Vasc Biol. 1997;17(6):1143–1143.

10. Nashel DJ. Is atherosclerosis a complication of long-term corticosteroid treatment? Am J Med. 1986;80(5):925–925.

11. MacLeod C, Hadoke PWF, and Nixon M. Glucocorticoids: Fuelling the Fire of Atherosclerosis or Therapeutic Extinguishers? Int J Mol Sci. 2021;22(14).

12. Pujades-Rodriguez M, Morgan AW, Cubbon RM, and Wu J. Dose-dependent oral glucocorticoid cardiovascular risks in people with immune-mediated in?ammatory diseases: A population-based cohort study. PLoS Med. 2020;17(12):e1003432.

13. Trusca VG, Fuior EV, Fenyo IM, Kardassis D, Simionescu M, and Gafencu AV. Differential action of glucocorticoids on apolipoprotein E gene expression in macrophages and hepatocytes. PloS one. 2017;12(3):e0174078.

14. Ayaori M, Sawada S, Yonemura A, Iwamoto N, Ogura M, Tanaka N, et al. Glucocorticoid receptor regulates ATP-binding cassette transporter-A1 expression and apolipoprotein-mediated cholesterol e?ux from macrophages. Arterioscler Thromb Vasc Biol. 2006;26(1):163–163.

15. Cavenee WK, Johnston D, and Melnykovych G. Regulation of cholesterol biosynthesis in HeLa S3G cells by serum lipoproteins: dexamethasone-mediated interference with suppression of 3-hydroxy-3-methylglutaryl coenzyme A reductase. Proc Natl Acad Sci U S A. 1978;75(5):2103–2103.

16. van Rossum EF, and Lamberts SW. Polymorphisms in the glucocorticoid receptor gene and their associations with metabolic parameters and body composition. Recent Prog Horm Res. 2004;59:333–57.

17. Yudt MR, and Cidlowski JA. The glucocorticoid receptor: coding a diversity of proteins and responses through a single gene. Mol Endocrinol. 2002;16(8):1719–1719.

18. van Rossum EF, Koper JW, Huizenga NA, Uitterlinden AG, Janssen JA, Brinkmann AO, et al. A polymorphism in the glucocorticoid receptor gene, which decreases sensitivity to glucocorticoids in vivo, is associated with low insulin and cholesterol levels. Diabetes. 2002;51(10):3128–3128.

19. Kino T. Single NucleoMde Variations of the Human GR Gene Manifested as Pathologic Mutations or Polymorphisms. Endocrinology. 2018;159(7):2506–2506.

20. van Rossum EF, Feelders RA, van den Beld AW, Uitterlinden AG, Janssen JA, Ester W, et al. Association of the ER22/23EK polymorphism in the glucocorticoid receptor gene with survival and C-reactive protein levels in elderly men. Am J Med. 2004;117(3):158–158.

21. Hu W, Jiang C, Kim M, Yang W, Zhu K, Guan D, et al. Individual-specific functional epigenomics reveals genetic determinants of adverse metabolic effects of glucocorticoids. Cell Metab. 2021;33(8):1592–1592 e7.

22. Koper JW, Stolk RP, de Lange P, Huizenga NA, Molijn GJ, Pols HA, et al. Lack of association between five polymorphisms in the human glucocorticoid receptor gene and glucocorticoid resistance. Hum Genet. 1997;99(5):663–663.

23. Saadatagah S, Jose M, Dikilitas O, Alhalabi L, Miller AA, Fan X, et al. Author Correction: Genetic basis of hypercholesterolemia in adults. NPJ Genom Med. 2021;6(1):56.

24. Trapani L, Segatto M, and Pallomni V. Regulation and deregulation of cholesterol homeostasis: The liver as a metabolic “power station”. World J Hepatol. 2012;4(6):184–184.

25. The Genotype-Tissue Expression (GTEx) project. Nat Genet. 2013;45(6):580–580.

26. Weikum ER, Knuesel MT, Ortlund EA, and Yamamoto KR. Glucocorticoid receptor control of transcription: precision and plasticity via allostery. Nat Rev Mol Cell Biol. 2017;18(3):159–159.

27. Russo P, Tomino C, Santoro A, Prinzi G, Proiem S, Kisialiou A, et al. FKBP5 rs4713916: A Potential Genetic Predictor of Interindividual Different Response to Inhaled Corticosteroids in Patients with Chronic Obstructive Pulmonary Disease in a Real-Life Semng. Int J Mol Sci. 2019;20(8).

28. Zannas AS, Wiechmann T, Gassen NC, and Binder EB. Gene-Stress-Epigenetic Regulation of FKBP5: Clinical and Translational Implications. Neuropsychopharmacology. 2016;41(1):261–261.

29. Hollstein T, Vogt A, Grenkowitz T, Stojakovic T, März W, Laufs U, et al. Treatment with PCSK9 inhibitors reduces atherogenic VLDL remnants in a real-world study. Vascul Pharmacol. 2019;116:8–15.

30. Canuel M, Sun X, Asselin MC, Paramithiotis E, Prat A, and Seidah NG. Proprotein convertase subtilisin/kexin type 9 (PCSK9) can mediate degradation of the low density lipoprotein receptor-related protein 1 (LRP-1). PloS one. 2013;8(5):e64145.

31. Maxwell KN, Fisher EA, and Breslow JL. Overexpression of PCSK9 accelerates the degradation of the LDLR in a post-endoplasmic reticulum compartment. Proc Natl Acad Sci U S A. 2005;102(6):2069–2069.

32. Lagace TA. PCSK9 and LDLR degradation: regulatory mechanisms in circulation and in cells. Curr Opin Lipidol. 2014;25(5):387–387.

33. Azmi S, Sun H, Ozog A, and Taneja R. mSharp-1/DEC2, a basic helix-loop-helix protein functions as a transcriptional repressor of E box activity and Stra13 expression. J Biol Chem. 2003;278(22):20098–20098.

34. Honma S, Kawamoto T, Takagi Y, Fujimoto K, Sato F, Noshiro M, et al. Dec1 and Dec2 are regulators of the mammalian molecular clock. Nature. 2002;419(6909):841–841.

35. Rouillard AD, Gundersen GW, Fernandez NF, Wang Z, Monteiro CD, McDermott MG, et al. The harmonizome: a collection of processed datasets gathered to serve and mine knowledge about genes and proteins. Database (Oxford). 2016;2016.

36. Shen WJ, Azhar S, and Kraemer FB. SR-B1: A Unique Multifunctional Receptor for Cholesterol In?ux and E?ux. Annu Rev Physiol. 2018;80:95–116.

37. Hamilton KA, Wang Y, Raefsky SM, Berkowitz S, Spangler R, Suire CN, et al. Mice lacking the transcriptional regulator Bhlhe40 have enhanced neuronal excitability and impaired synaptic plasticity in the hippocampus. PloS one. 2018;13(5):e0196223.

38. Peters DT, Henderson CA, Warren CR, Friesen M, Xia F, Becker CE, et al. Asialoglycoprotein receptor 1 is a specific cell-surface marker for isolating hepatocytes derived from human pluripotent stem cells. Development. 2016;143(9):1475–1475.

39. Lu H, Lei X, Winkler R, John S, Kumar D, Li W, et al. Crosstalk of hepatocyte nuclear factor 4a and glucocorticoid receptor in the regulation of lipid metabolism in mice fed a high-fat-high-sugar diet. Lipids Health Dis. 2022;21(1):46.

40. Oyadomari S, Matsuno F, Chowdhury S, Kimura T, Iwase K, Araki E, et al. The gene for hepatocyte nuclear factor (HNF)-4alpha is activated by glucocorticoids and glucagon, and repressed by insulin in rat liver. FEBS LeP. 2000;478(1-2):141–6.

41. Engblom D, Kornfeld JW, Schwake L, Tronche F, Reimann A, Beug H, et al. Direct glucocorticoid receptor-Stat5 interaction in hepatocytes controls body size and maturation-related gene expression. Genes Dev. 2007;21(10):1157–1157.

42. Bharathan SP, Manian KV, Aalam SM, Palani D, Deshpande PA, Pratheesh MD, et al. Systematic evaluation of markers used for the identification of human induced pluripotent stem cells. Biol Open. 2017;6(1):100–100.

43. Wang P, McKnight KD, Wong DJ, Rodriguez RT, Sugiyama T, Gu X, et al. A molecular signature for purified definitive endoderm guides di?erentiation and isolation of endoderm from mouse and human embryonic stem cells. Stem Cells Dev. 2012;21(12):2273–2273.

44. Ghosheh N, Olsson B, Edsbagge J, Kuppers-Munther B, Van Giezen M, Asplund A, et al. Highly Synchronized Expression of Lineage-Specific Genes during In Vitro Hepatic Differentiation of Human Pluripotent Stem Cell Lines. Stem Cells Int. 2016;2016:8648356.

45. Siller R, Greenhough S, Naumovska E, and Sullivan GJ. Small-molecule-driven hepatocyte di?erentiation of human pluripotent stem cells. Stem Cell Reports. 2015;4(5):939–939.

46. de Villiers WJ, van der Westhuyzen DR, Coetzee GA, Henderson HE, and Marais AD. The apolipoprotein E2 (Arg145Cys) mutation causes autosomal dominant type III hyperlipoproteinemia with incomplete penetrance. Arterioscler Thromb Vasc Biol. 1997;17(5):865–865.

47. Sullivan PM, Mezdour H, Quarfordt SH, and Maeda N. Type III hyperlipoproteinemia and spontaneous atherosclerosis in mice resulting from gene replacement of mouse Apoe with human Apoe*2. J Clin Invest. 1998;102(1):130–130.

48. Huang Y, Schwendner SW, Rall SC, Jr., and Mahley RW. Hypolipidemic and hyperlipidemic phenotypes in transgenic mice expressing human apolipoprotein E2. J Biol Chem. 1996;271(46):29146–29146.

49. Backstrom JR, Sheng J, Wang MC, Bernardo-Colón A, and Rex TS. Optimization of S. aureus dCas9 and CRISPRi Elements for a Single Adeno-Associated Virus that Targets an Endogenous Gene. Mol Ther Methods Clin Dev. 2020;19:139–48.

50. Malisch JL, Saltzman W, Gomes FR, Rezende EL, Jeske DR, and Garland T, Jr. Baseline and stress-induced plasma corticosterone concentrations of mice selectively bred for high voluntary wheel running. Physiol Biochem Zool. 2007;80(1):146–146.

51. Hatch N, Srodulski S, Chan H-W, Zhang X, Tannock L, and King V. Endogenous Androgen Deficiency Enhances Diet-Induced Hypercholesterolemia and Atherosclerosis in Low-Density Lipoprotein Receptor-Deficient Mice. Gender medicine. 2012;9:319–28.

52. Lei Z, Wu H, Yang Y, Hu Q, Lei Y, Liu W, et al. Ovariectomy Impaired Hepatic Glucose and Lipid Homeostasis and Altered the Gut Microbiota in Mice With Different Diets. Front Endocrinol (Lausanne). 2021;12:708838.

53. McCracken KW, Catá EM, Crawford CM, Sinagoga KL, Schumacher M, Rockich BE, et al. Modelling human development and disease in pluripotent stem-cell-derived gastric organoids. Nature. 2014;516(7531):400–400.

54. Schmidt A, Harada SI, Kimmel DB, Bai C, Chen F, Rutledge SJ, et al. Identification of anabolic selective androgen receptor modulators with reduced activities in reproductive tissues and sebaceous glands. J Biol Chem. 2009;284(52):36367–36367.

55. Brovkina AF, Sychev DA, and Toropova OS. [In?uence of CYP3A4, CYP3A5, and NR3C1 genes polymorphism on the e?ectiveness of glucocorticoid therapy in patients with endocrine ophthalmopathy]. Vestn ORalmol. 2020;136(6. Vyp. 2):125–32.

56. Constantinescu AE, Mitchell RE, Zheng J, Bull CJ, Timpson NJ, Amulic B, et al. A framework for research into continental ancestry groups of the UK Biobank. Hum Genomics. 2022;16(1):3.

57. Russcher H, Smit P, van den Akker EL, van Rossum EF, Brinkmann AO, de Jong FH, et al. Two polymorphisms in the glucocorticoid receptor gene directly affect glucocorticoid-regulated gene expression. J Clin Endocrinol Metab. 2005;90(10):5804–5804.

58. El-Fayoumi R, Hagras M, Abozenadaha A, Bawazir W, and Shinawi T. Association Between NR3C1 Gene Polymorphisms and Toxicity Induced by Glucocorticoids Therapy in Saudi Children with Acute Lymphoblastic Leukemia. Asian Pac J Cancer Prev. 2018;19(5):1415–1415.

59. Roerink SH, Wagenmakers MA, Smit JW, van Rossum EF, Netea-Maier RT, Plantinga TS, et al. Glucocorticoid receptor polymorphisms modulate cardiometabolic risk factors in patients in long-term remission of Cushing’s syndrome. Endocrine. 2016;53(1):63–63.

60. Quax RA, Koper JW, Huisman AM, Weel A, Hazes JM, Lamberts SW, et al. Polymorphisms in the glucocorticoid receptor gene and in the glucocorticoid-induced transcript 1 gene are associated with disease activity and response to glucocorticoid bridging therapy in rheumatoid arthritis. Rheumatol Int. 2015;35(8):1325–1325.

61. Bouma EM, Riese H, Nolte IM, Oosterom E, Verhulst FC, Ormel J, et al. No associations between single nucleotide polymorphisms in corticoid receptor genes and heart rate and cortisol responses to a standardized social stress test in adolescents: the TRAILS study. Behav Genet. 2011;41(2):253–253.

62. Larsson CA, Gullberg B, Råstam L, and Lindblad U. Salivary cortisol differs with age and sex and shows inverse associations with WHR in Swedish women: a cross-sectional study. BMC Endocr Disord. 2009;9:16.

63. Durumutla HB, Prabakaran A, El Abdellaoui Soussi F, Akinborewa O, Latimer H, McFarland K, et al. Glucocorticoid chrono-pharmacology promotes glucose metabolism in heart through a cardiomyocyte-autonomous transactivation program. JCI Insight. 2024.

64. Prabakaran AD, McFarland K, Miz K, Durumutla HB, Piczer K, El Abdellaoui Soussi F, et al. Intermittent glucocorticoid treatment improves muscle metabolism via the PGC1alpha/Lipin1 axis in an aging-related sarcopenia model. J Clin Invest. 2024;134(11).

65. Chang CC, Chow CC, Tellier LC, Vamkuti S, Purcell SM, and Lee JJ. Second-generation PLINK: rising to the challenge of larger and richer datasets. Gigascience. 2015;4:7.

