## Supplemental Material bundle for "The human glucocorticoid receptor variant rs6190 increases blood cholesterol and promotes atherosclerosis"

### Supplementary Figure 1

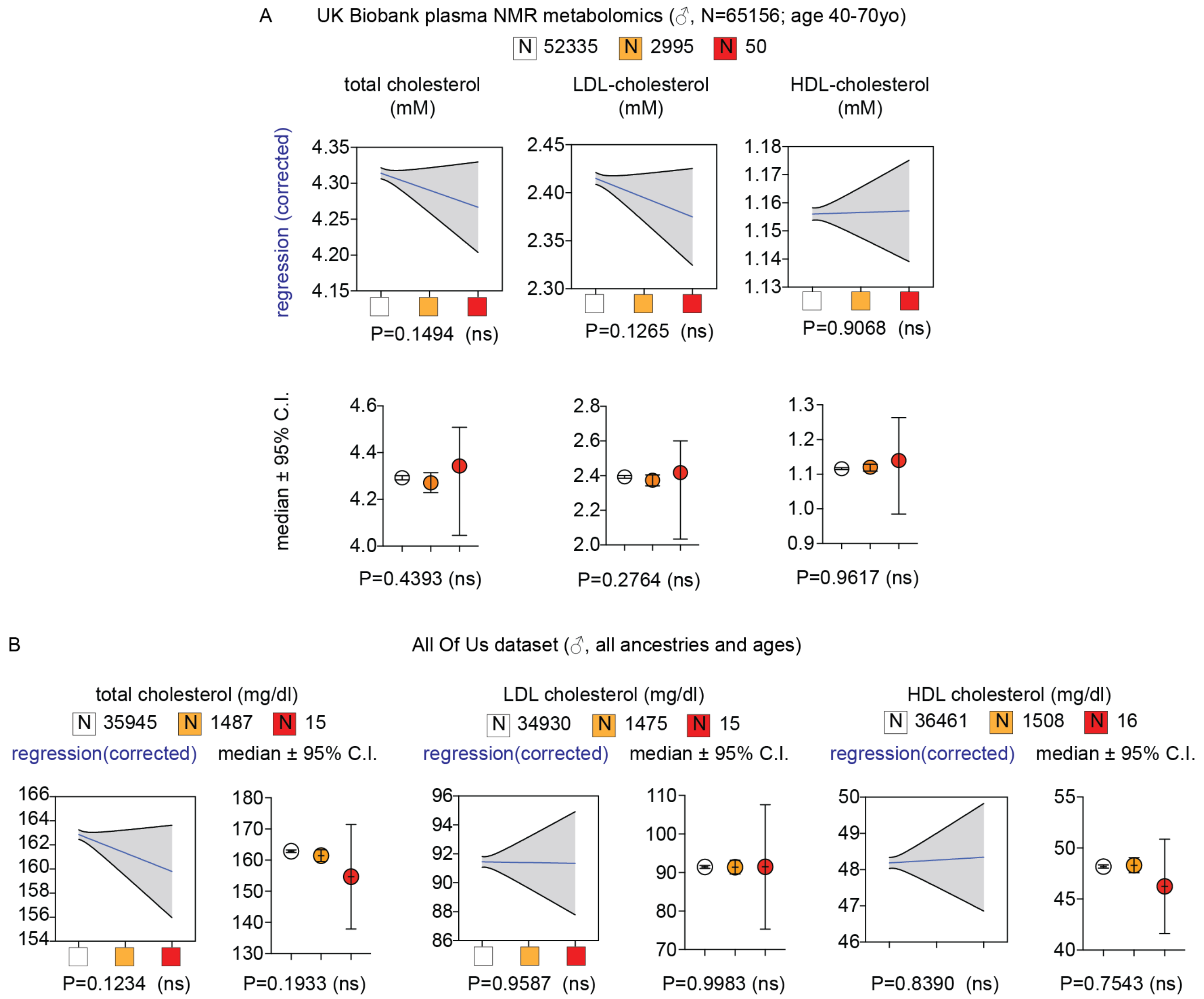


**Supplementary Figure 1. Related to Figure 1. Additional data from UK Biobank and All Of Us datasets.** Zygosity-dependent correlations of rs6190 with total, LDL and HDL cholesterol were not significant in men from either UK Biobank (A) or All of Us (B) datasets. Linear regressions were corrected for age, diabetes, triacylglycerols; median intervals were compared through Kruskal-Wallis test.

#### Supplementary Figure 2

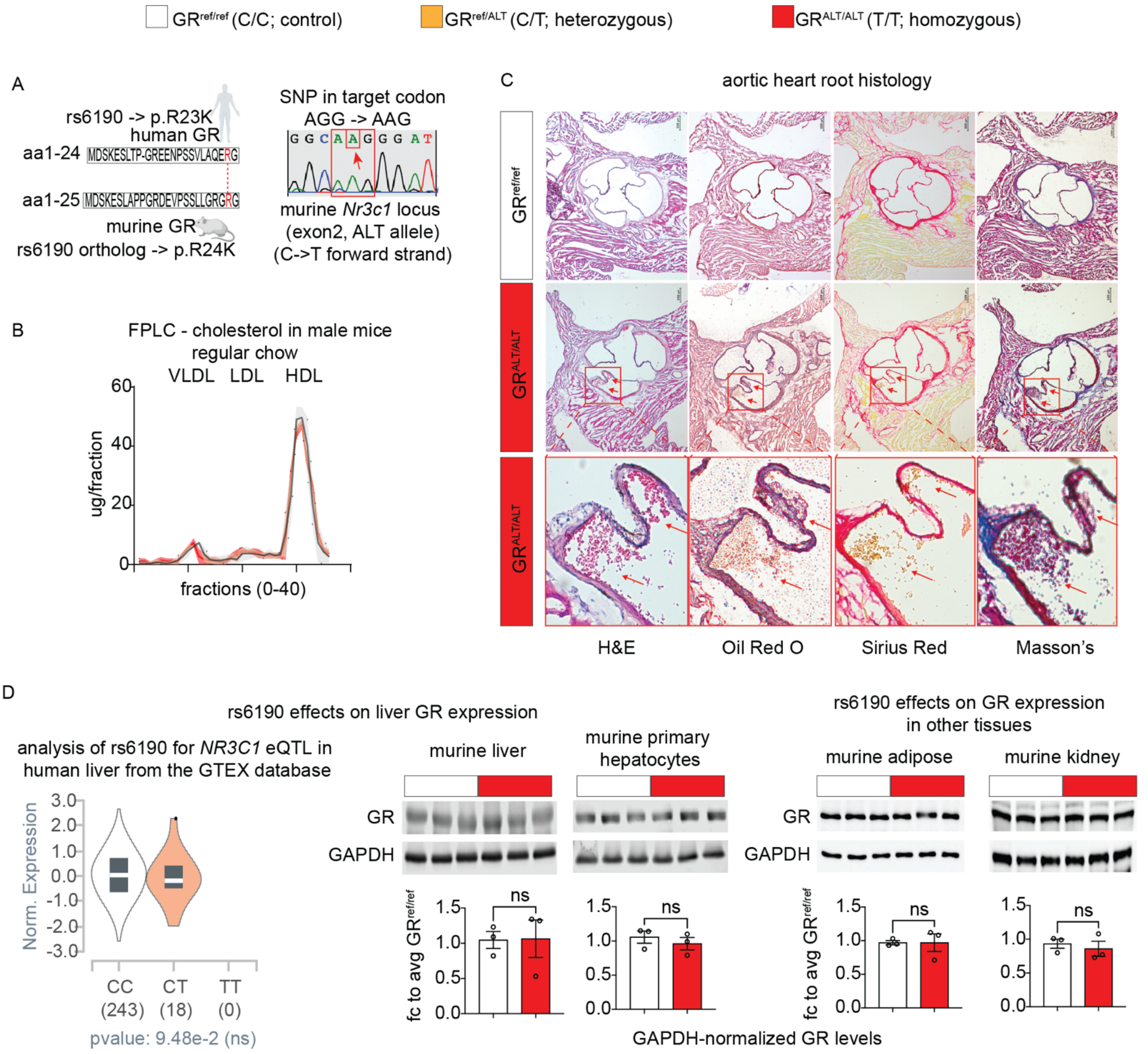


**Supplementary Figure 2. Related to Figure 2. Additional analyses related to mutant GR effects in murine liver. (A)** Diagram highlighting the human-mouse GR sequence orthology and the SNP genocopy introduced through CRISPR-Cas9. **(B)** Differently than in female mice, male mice blunted the SNP effect on cholesterol elevation according to SNP zygosity, according to FPLC cholesterol levels across lipoprotein fractions. **(C)** In 3 out of 5 female mice analyzed from the parental WT genetic background after Western diet exposure, emerging immature plaques were noted in the aortic roots (arrows; insets: high magnification). **(D)** Analysis of GTEX database and GR protein levels in livers, primary hepatocytes, white adipose tissue (ventral fat pad) or kidney of $GR^{ref/ref}$ vs $GR^{ALT/ALT}$ mice revealed no sizable effect of rs6190 on hepatic GR expression and protein levels. Scale bars, 100 µm. N=3-5/group, ♂ in B, ♀ in C-D, 6mo; D: Welch's t-test; ns, not significant.

#### Supplementary Figure 3

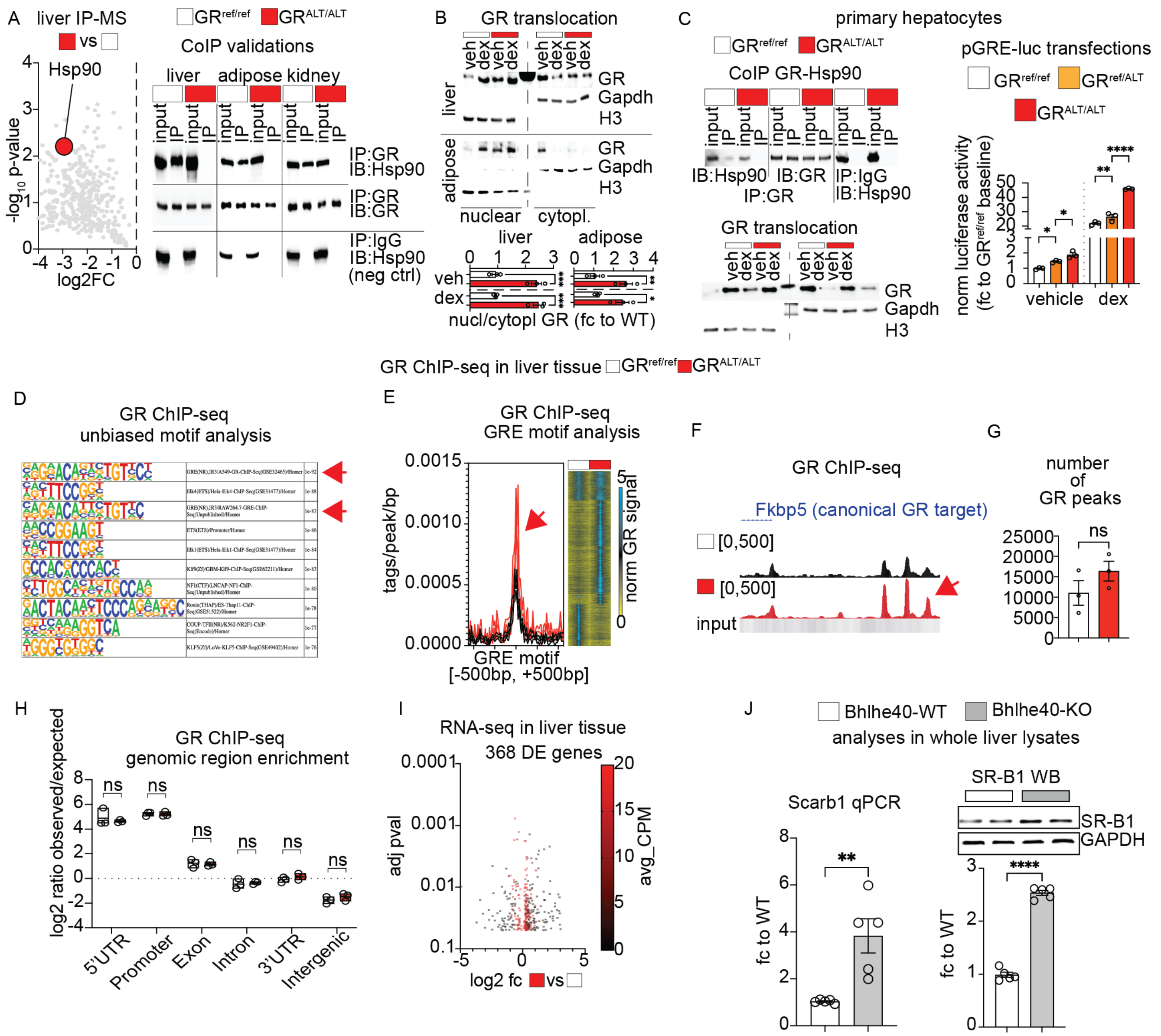


**Supplementary Figure 3. Related to Figure 2. Additional analyses related to mutant GR epigenetic regulation. (A)** IP-MS in liver identified Hsp90 among top hits for decreased interaction with the mutant GR compared to the WT GR. CoIP in tissue extracts confirmed decreased GR-Hsp90 interaction in not only liver, but also other metabolically active tissues like adipose and kidney. **(B)** At 30min after one i.p. 1mg/kg dexamethasone (dex) or vehicle (veh), the mutant GR showed increased nuclear/cytoplasmic signal enrichment compared to WT GR in liver, adipose and kidney. H3, nuclear fractions; Gapdh, cytoplasmic fractions. **(C)** Primary hepatocytes recapitulated the mutant vs WT GR differences in Hsp90 interaction and nuclear enrichment. Moreover, the SNP increased basal and steroid-driven GR activity on a GRE luciferase reporter transfected in primary hepatocytes according to SNP zygosity. **(D)** Unbiased motif analysis validated ChIP-seq datasets through enrichment for GRE motif (arrows). **(E)** Mutant GR showed increased GR occupancy genome-wide on GRE motifs (arrow). **(F)** Representative peak traces for a canonical marker of GR epigenetic activity, the *Fkbp5* distal promoter, showed increased GR occupancy on canonical GR sites (arrows). **(G-H)** Total number of peaks did not significantly increase for the mutant GR compared to the control GR, and the relative distribution of peaks into gene loci regions did not change. **(I)** Volcano plot of SNP-dependent differentially expressed genes in liver per RNA-seq datasets. **(J)** Compared to *Bhlhe40-WT* littermates, *Bhlhe40-KO* livers showed upregulation of SR-B1 levels at mRNA (*Scarb1*, gene name for SR-B1), supporting the notion of Bhlhe40 as transcriptional repressor of SR-B1 in liver. N=3-5/group, ♂ in B, ♀ in C-L, 6mo; D, E, J: 2w ANOVA + Sidak; I, L: Welch's t-test; *, $P<0.05$; **, $P<0.01$; ***, $P<0.001$; ****, $P<0.0001$.

#### Supplementary Figure 4

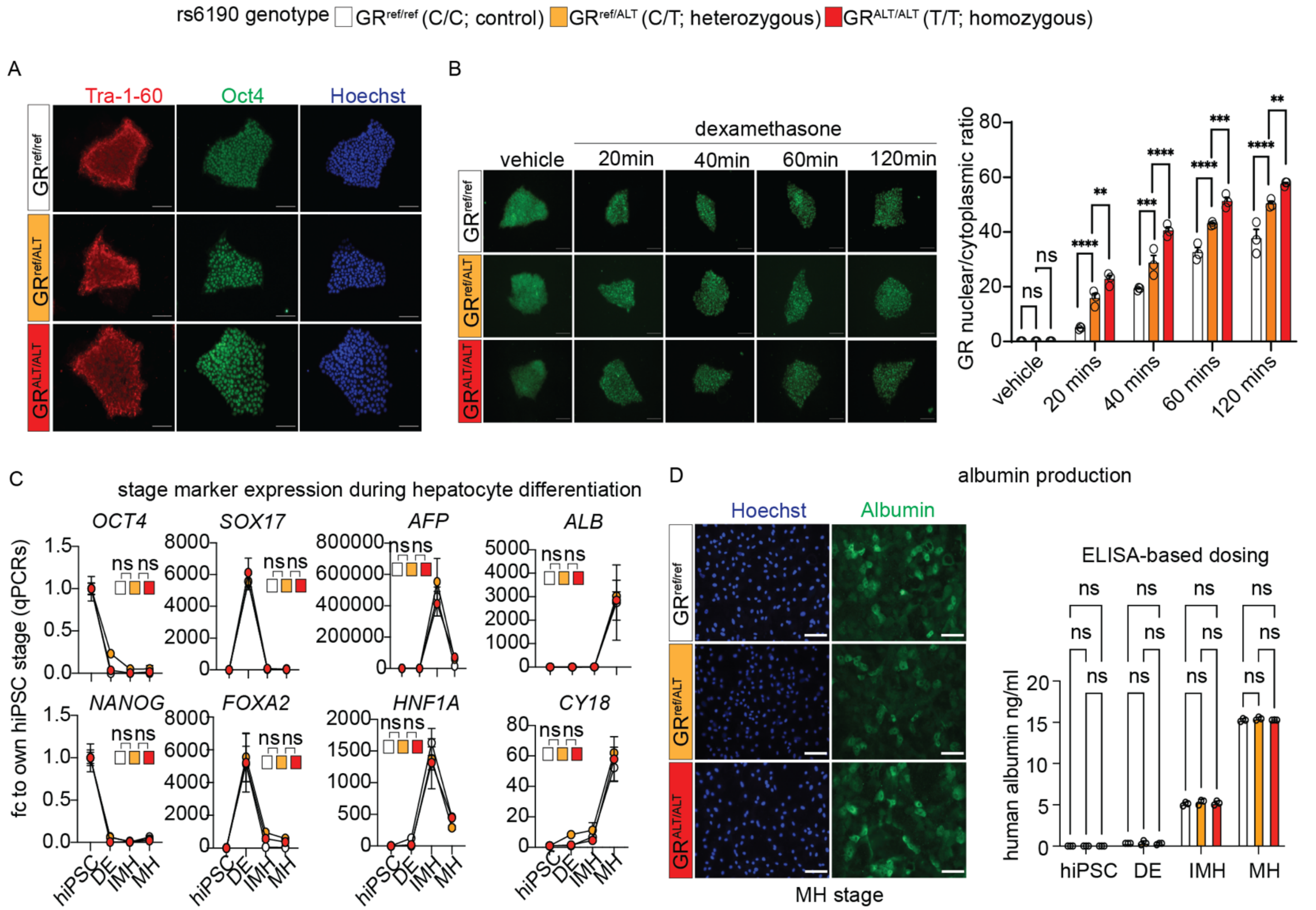


**Supplementary Figure 4. Related to Figure 3. Additional analyses related to SNP-mutant hiPSCs. (A)** Pluripotency marker validation of isogenic hiPSC lines with CRISPR-knock-in engineering of a SNP genocopy in the endogenous *NR3C1* gene locus. **(B)** The SNP promoted GR translocation at the undifferentiated hiPSC stage. **(C)** The SNP genotype did not impact the overall progression of differentiating hiPSCs across the stages of hepatocyte differentiation: hiPSC, pluripotent; DE, definitive endoderm; IMH, immature hepatocytes; MH, mature hepatocytes. (**D**) Albumin production (staining) and secretion (ELISA) confirmed hepatocyte maturation comparable across SNP genotypes. Scale bars, 100 μm. Each dot represents an independent differentiation replicate; N=3/group. 2w ANOVA + Sidak; *, P<0.05; **, P<0.01; ***, P<0.001; ****, P<0.0001.

**Supplementary Figure 5**

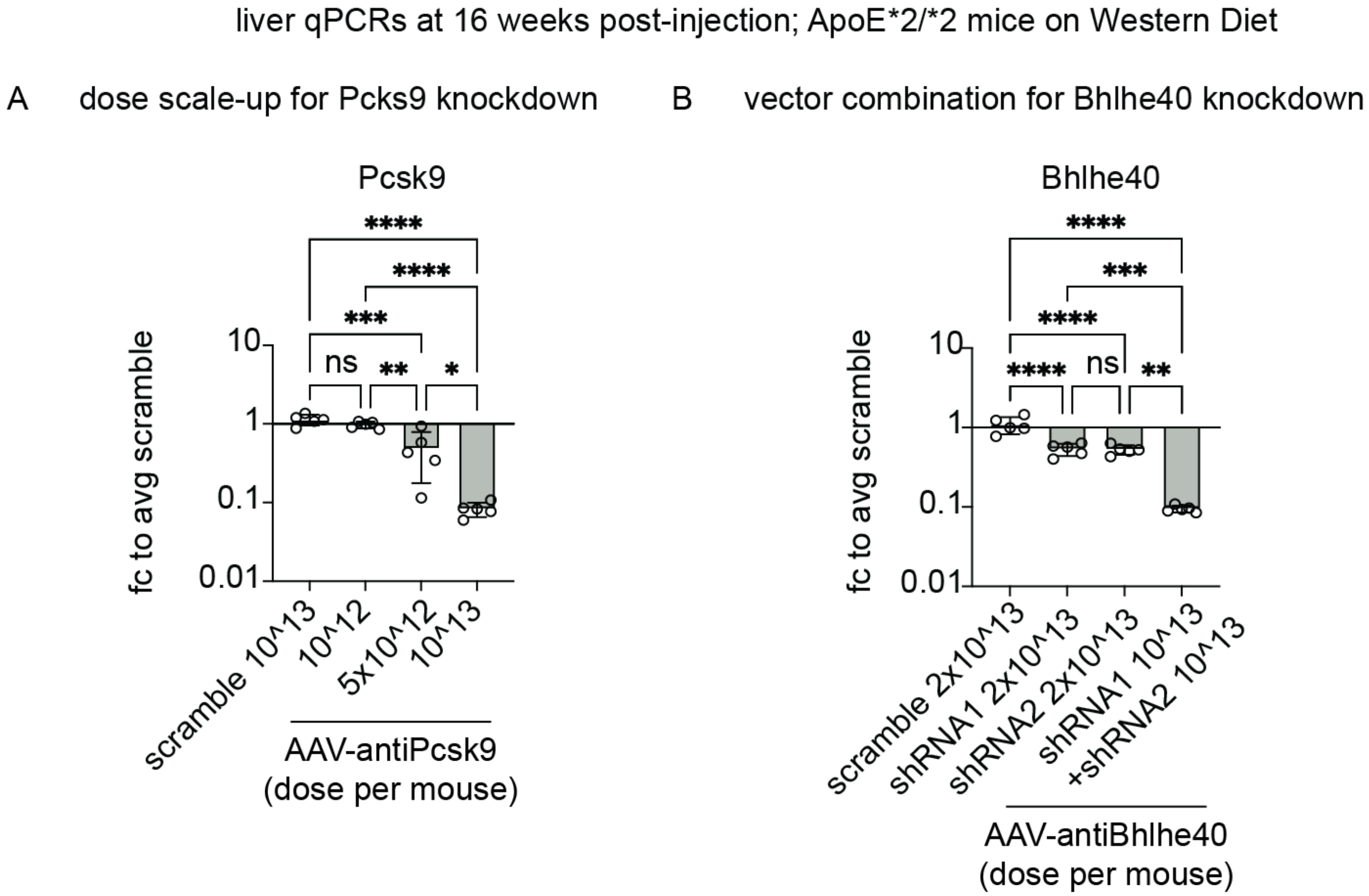


**Supplementary Figure 5 – Preliminary validations of dosage and combinations for AAV8-mediated knockdowns in vivo. (A)** 10^13vg/mouse maximized *Pcsk9* knockdown in liver with AAV8-antiPcsk9 compared to scramble. **(B)** Combination of both AAV8-antiBhlhe40 shRNA vectors was synergistic in maximizing *Bhlhe40* knockdown in liver compared to scramble. N=5♀/group, 6mo; 1w ANOVA + Sidak; *, $P<0.05$; **, $P<0.01$; ***, $P<0.001$; ****, $P<0.0001$.

#### Supplementary Table 1

| | | | EBI GWAS Catalog | | | | | All of Us - sex-disaggregated regressions for total cholesterol | | | | | | |
|---|---|---|---|---|---|---|---|---|---|---|---|---|---|---|
| MAPPED_GENE | variant | location per dbSNP | P-VALUE | DISEASE/TRAIT | PUBMEDID | FIRST AUTHOR | DATE | N of non-carriers | N of heterozygous carriers | N of homozygous carriers | beta (men) | adj p-value (men) | beta (women) | adj p-value (women) |
| NR3C1 | rs10482682 | intron | 1E-60 | Height | 36224396 | Yengo L | 10/12/22 | 202223 | 168777 | 43810 | 0.098629 | 0.63895 | 1.854685 | **2.94958E-11** |
| NR3C1 | rs10482714 | 3' UTR | 2E-33 | Height | 36224396 | Yengo L | 10/12/22 | 409144 | 5646 | 24 | 0.278575 | 0.804081 | 2.903106 | 0.1214012 |
| NR3C1 | rs11747997 | intron | 1E-13 | Height | 34594039 | Sakaue S | 9/30/21 | 215896 | 163915 | 34329 | -0.344339 | 0.11624 | 0.864988 | 0.06155634 |
| NR3C1 | rs11750560 | intergenic | 7E-14 | Lung function (FVC) | 30595370 | Kichaev G | 12/27/18 | 103220 | 202825 | 108767 | -0.165463 | 0.407181 | 0.332345 | 0.563788 |
| NR3C1 | rs12187548 | intergenic | 0.00000001 | Body mass index (MTAG) | 36376304 | Koskeridis F | 11/14/22 | 476449 | 177787 | 175074 | 0.101692 | 0.417897 | -0.397485 | 0.05615418 |
| NR3C1 | rs13361249 | intergenic | 0.00000002 | Protein quantitative trait loci (liver) | 32778093 | He B | 8/10/20 | 1236005 | 8204 | 68 | -1.775785 | 0.085385 | -3.117363 | **0.000394715** |
| NR3C1 | rs142193461 | intergenic | 2E-13 | Bone mineral density mean | 37500982 | He D | 7/27/23 | 824237 | 5350 | 25 | 0.189691 | 0.875655 | 2.152551 | 0.1212883 |
| NR3C1 | rs1635474 | intergenic | 3E-11 | Hemoglobin | 32888494 | Vuckovic D | 9/1/20 | 215176 | 164861 | 34770 | -0.299896 | 0.170599 | 0.7605 | 0.07968305 |
| | | | 2E-10 | Hemoglobin concentration | 32888493 | Chen MH | 9/1/20 | | | | | | | |
| NR3C1 | rs17100350 | intergenic | 6E-11 | Height | 36224396 | Yengo L | 10/12/22 | 295183 | 96701 | 22931 | -0.350268 | 0.182487 | -4.014213 | **3.5059E-41** |
| NR3C1 | rs17287745 | intergenic | 0.00000004 | percentage (adjusted for testosterone an | 37867527 | Roshandel D | 10/5/23 | 604786 | 173998 | 50840 | -0.026703 | 0.872887 | 1.291901 | **5.57387E-08** |
| NR3C1 | rs17287758 | 3' UTR | 0.00000002 | Free testosterone levels | 36653534 | Leinonen JT | 1/18/23 | 308945 | 97116 | 8754 | -0.318322 | 0.255951 | 1.413211 | **0.006045405** |
| NR3C1 | rs174047 | intergenic | 3E-14 | Height | 39134668 | Shi S | 8/12/24 | 106549 | 205505 | 102715 | -0.318461 | 0.11132 | 0.784693 | 0.061568 |
| NR3C1 | rs174048 | intergenic | 1E-11 | Atrial fibrillation | 29892015 | Roselli C | 6/11/18 | 729897 | 92132 | 7583 | -0.282572 | 0.304529 | 0.504352 | 0.3634234 |
| | | | 6E-11 | Atrial fibrillation | 29892015 | Roselli C | 6/11/18 | | | | | | | |
| NR3C1 | rs2121152 | intergenic | 0.00000002 | tension (confirmatory factor analysis Fac | 38965376 | Carey CE | 7/4/24 | 460576 | 178542 | 190530 | 0.057521 | 0.634487 | 0.396697 | **0.003277398** |
| NR3C1 | rs2398629 | intron | 7E-13 | Lymphocyte percentage of white cells | 32888494 | Vuckovic D | 9/1/20 | 308390 | 97548 | 8853 | -0.317045 | 0.256827 | 1.408794 | **0.006389321** |
| | | | 2E-11 | Neutrophil percentage of white cells | 32888494 | Vuckovic D | 9/1/20 | | | | | | | |
| | | | 1E-10 | mphocytes in blood (confirmatory factor a | 38965376 | Carey CE | 7/4/24 | | | | | | | |
| NR3C1 | rs258753 | intergenic | 7E-14 | Hematocrit | 32888493 | Chen MH | 9/1/20 | 221386 | 161009 | 32400 | -0.309872 | 0.16066 | 0.892954 | **0.03415907** |
| | | | 1E-12 | Hemoglobin concentration | 32888493 | Chen MH | 9/1/20 | | | | | | | |
| | | | 3E-11 | Hematocrit | 32888493 | Chen MH | 9/1/20 | | | | | | | |
| | | | 4E-11 | Hematocrit | 32888494 | Vuckovic D | 9/1/20 | | | | | | | |
| NR3C1 | rs258755 | intron | 0.00000005 | Height (standard GWA) | 37106081 | Schoeler T | 4/27/23 | 218167 | 163055 | 33565 | -0.293575 | 0.182271 | 0.762332 | 0.06973425 |
| NR3C1 | rs258756 | intergenic | 0.000000003 | Eosinophil percentage of white cells | 32888494 | Vuckovic D | 9/1/20 | 109750 | 200884 | 104063 | -0.198004 | 0.317794 | 1.599138 | **1.36503E-08** |
| | | | 0.000000006 | Eosinophil counts | 32888494 | Vuckovic D | 9/1/20 | | | | | | | |
| NR3C1 | rs258762 | intergenic | 0.00000004 | Multi-trait sex score | 37277458 | Vosberg DE | 6/5/23 | 2591742 | 204555 | 104209 | -0.03576 | 0.763476 | 0.714528 | **1.69901E-05** |
| NR3C1 | rs258763 | intergenic | 1E-15 | Lung function (forced vital capacity) | 36914875 | Shrine N | 3/13/23 | 534263 | 198103 | 97268 | -0.084042 | 0.553139 | 0.9182 | **0.00001** |
| | | | 0.00000005 | Hip circumference adjusted for BMI | 34021172 | Christakoudi S | 5/21/21 | | | | | | | |
| NR3C1 | rs258796 | intergenic | 0.000000001 | Eosinophil counts | 30595370 | Kichaev G | 12/27/18 | 231301 | 154736 | 28752 | -0.323557 | 0.150457 | 1.576241 | **7.79509E-06** |
| NR3C1 | rs28674017 | intergenic | 0.00000005 | Medication use (thyroid preparations) | 31015401 | Wu Y | 4/23/19 | 222942 | 161658 | 30203 | 0.466019 | 0.038129 | 1.173322 | **2.59967E-08** |
| NR3C1 | rs34632394 | intergenic | 3E-10 | Total testosterone levels | 32042192 | Ruth KS | 2/10/20 | 145416 | 189045 | 80328 | 0.380686 | 0.053105 | 2.167328 | **8.38434E-22** |
| | | | 9E-10 | Total testosterone levels | 32042192 | Ruth KS | 2/10/20 | | | | | | | |
| NR3C1 | rs4912652 | intergenic | 1E-10 | Body mass index | 36581621 | Huang J | 12/29/22 | 479759 | 187331 | 162544 | 0.101525 | 0.428854 | -0.408897 | 0.06699651 |
| NR3C1 | rs6198 | 3' UTR | 9E-15 | Height | 36224396 | Yengo L | 10/12/22 | 316484 | 90094 | 8178 | -0.392892 | 0.168458 | 1.982386 | **0.000044** |
| NR3C1 | rs6580277 | intergenic | 2E-19 | Atrial fibrillation | 36653681 | Miyazawa K | 1/19/23 | 263563 | 132173 | 18999 | 0.075197 | 0.758628 | 0.201054 | 0.5304 |
| | | | 2E-17 | Atrial fibrillation | 30061737 | Nielsen JB | 7/30/18 | | | | | | | |
| | | | 7E-17 | Atrial fibrillation (MTAG) | 39537608 | Koskeridis F | 11/13/24 | | | | | | | |
| | | | 1E-16 | Atrial fibrillation (MTAG) | 35872910 | Carcel-Marquez J | 7/8/22 | | | | | | | |
| NR3C1 | rs6860760 | intergenic | 0.00000003 | Metabolic syndrome | 39349817 | Park S | 9/30/24 | 480184 | 187508 | 161868 | 0.092357 | 0.472223 | -0.429333 | **0.04078205** |
| NR3C1 | rs6865292 | intron | 0.000000008 | Free androgen index | 36653534 | Leinonen JT | 1/18/23 | 681614 | 128426 | 19558 | -0.155758 | 0.467206 | 1.182242 | **0.001656039** |
| NR3C1 | rs6877893 | intron | 0.000000009 | t circumference adjusted for body mass | 31669095 | Zhu Z | 10/24/19 | 96424 | 202622 | 115771 | 0.061278 | 0.759042 | -1.157755 | **3.73439E-05** |
| NR3C1 | rs72801051 | intron | 0.00000004 | Carotid intima media thickness (mean) | 34852643 | Wai Yeung M | 12/2/21 | 747119 | 74097 | 5862 | -0.292621 | 0.324435 | 1.748319 | **0.000291134** |
| NR3C1 | rs72802806 | intron | 0.000000009 | rmone-binding globulin levels adjusted f | 32042192 | Ruth KS | 2/10/20 | 298212 | 104420 | 12167 | -0.28367 | 0.286403 | 1.470548 | **0.003016232** |
| | | | 0.00000002 | Sex hormone-binding globulin levels | 36653534 | Leinonen JT | 1/18/23 | | | | | | | |
| NR3C1 | rs72802813 | intron | 0.000000001 | Hip circumference adjusted for BMI | 34021172 | Christakoudi S | 5/21/21 | 286019 | 115646 | 13109 | 0.101162 | 0.695745 | 1.36654 | **1.15264E-05** |
| NR3C1 | rs74601840 | intergenic | 0.000000004 | anti-Xa activity of apixaban | 36867504 | Mu G | 3/3/23 | 413489 | 1064 | 20 | 10.036885 | 0.000682 | 2.615795 | **0.001919059** |
| NR3C1 | rs7701443 | intron | 0.000000006 | Hip circumference adjusted for BMI | 34021172 | Christakoudi S | 5/21/21 | 548150 | 202866 | 78596 | 0.15334 | 0.318124 | -0.216051 | 0.8199622 |
| NR3C1 | rs7724289 | intergenic | 3E-25 | Height | 36224396 | Yengo L | 10/12/22 | 326327 | 82413 | 6038 | 0.621465 | 0.051528 | -0.314938 | 0.4520151 |
| NR3C1 | rs852982 | intron | 1E-73 | Height | 36224396 | Yengo L | 10/12/22 | 635127 | 161554 | 32905 | -0.269435 | 0.148763 | 0.699489 | 0.07909547 |
| NR3C1 | rs853175 | intergenic | 2E-22 | Height | 30595370 | Kichaev G | 12/27/18 | 947703 | 198968 | 97765 | -0.073666 | 0.575111 | 0.765868 | **6.98636E-05** |
| | | | 1E-10 | Body surface area | 36502284 | Yu XH | 12/10/22 | | | | | | | |
| | | | 0.00000004 | Multi-trait sex score | 37277458 | Vosberg DE | 6/5/23 | | | | | | | |
| NR3C1 | rs864354 | intergenic | 9E-12 | Lung function (FEV1/FVC) | 30595370 | Kichaev G | 12/27/18 | 117897 | 198950 | 97955 | -0.173086 | 0.381023 | 1.693213 | **2.49153E-09** |

**Supplementary Table 1 – Additional analyses in NR3C1-related variants -** We extracted 38 *NR3C1* locus variants with GWAS significant hits (<5 × 10−8) from the current EBI GWAS catalog and ran linear regressions - aggregated for ancestry but disaggregated for sex - for total cholesterol in the All of Us dataset. All 38 variants were non-coding and in weak-to-negligible LD range with rs6190 (LD, $r^2$<0.15). None of the 38 variants had a significant reported GWAS hit for cholesterol. However, our regressions showed 24 variants with correlations (adj p-val <0.05; 20, direct; 4, inverse) with total cholesterol according to zygosity. Intriguingly, those correlations were significant only in women and not in men for all 24 variants, analogous to our findings with rs6190.

#### Supplementary Methods

##### Assays for circulating cholesterol, estradiol and testosterone

Blood samples were procured from ~3-month-old mice and collected in EDTA-treated tubes using cardiac puncture method following an overnight fasting. The blood samples were maintained on ice and subjected to centrifugation at 2500 x g for 10 mins to isolate plasma. Following the centrifugation step, the obtained plasma was immediately transferred into a clean microcentrifuge tube for plasma lipid measurements. The plasma levels of total cholesterol (TC) were measured using Infinity™ Cholesterol kit (Cat #TR13421, Thermo Fisher Scientific) and Infinity™ Triglyceride kit (Cat # TR22421, Thermo Fisher Scientific). The concentrations of estradiol (Cat # 501890, Cayman Chemicals) and testosterone (Cat #582701, Cayman Chemicals) in serum were measured according to the manufacturer's protocols for each kit.

##### RNA extraction and RT-qPCR

Total RNA was extracted from cryo-pulverized liver tissues and hiPSC-derived hepatocyte-like cells using Trizol (Cat #15596026, Thermo Fisher Scientific) and 1 ug RNA was reverse-transcribed using SuperScript™ IV VILO™ Master Mix (Cat #11756050, Thermo Fisher Scientific). RT-qPCRs were conducted in three replicates using 1X SYBR Green Fast qPCR machine (Bio-Rad, Hercules, CA; thermal profile: 95C, 15 sec; 60C, 30sec; 40x; melting curve). The 2-$\Delta\Delta$CT method was used to calculate relative gene expression. GAPDH was used as the internal control. Primers were selected among validated primer sets from MGH PrimerBank:

| Gene Name | Forward sequence | Reverse Sequence |
|---|---|---|
| Mouse *Pcsk9* | GAGACCCAGAGGCTACAGATT | AATGTACTCCACATGGGGCAA |
| Mouse *Bhlhe40* | ACGGAGACCTGTCAGGGATG | GGCAGTTTGTAAGTTTCCTTGC |
| Mouse *Scarb1* | TTTGGAGTGGTAGTAAAAAGGGC | TGACATCAGGGACTCAGAGTAG |
| Mouse *Ldlr* | TGACTCAGACGAACAAGGCTG | ATCTAGGCAATCTCGGTCTCC |
| Mouse *Gapdh* | GTATGACTCCACTCACGGCAAA | GGTCTCGCTCCTGGAAGATG |
| Human *OCT4* | AGCGAACCAGTATCGAGAAC | TTACAGAACCACACTCGGAC |

| | | |
|---|---|---|
| Human *NANOG* | CTCCAACATCCTGAACCTCAGC | CGTCACACCATTGCTATTCTTCG |
| Human *SOX17* | TATTTTGTCTGCCACTTGAACAGT | TTGGGACACATTCAAAGCTAGTTA |
| Human *FOXA2* | GCATTCCCAATCTTGACACGGTGA | GCCCTTGCAGCCAGAATACACATT |
| Human *NESTIN* | CTGCTACCCTTGAGACACCTG | GGGCTCTGATCTCTGCATCTAC |
| Human *PAX6* | AACGATAACATACCAAGCGTGT | GGTCTGCCCGTTCAACATC |
| Human *TBX6* | GTGTCTTTCCATCGTGTCAAGC | TATGCGGGGTTGGTACTTGTG |
| Human *MIXL1* | GGCGTCAGAGTGGGAAATCC | GGCAGGCAGTTCACATCTACC |
| Human *ALB* | CCCCAAGTGTCAACTCCA | GTTCAGGACCACGGATAG |
| Human *AFP* | ACTGAATCCAGAACACTGCA | TGCAGTCAATGCATCTTTCA |
| Human *HNF1A* | ACATGGACATGGCCGACTAC | CGTTGAGGTTGGTGCCTTCT |
| Human *CY18* | GCTGGAAGATGGCGAGGACTTT | TGGTCTCAGACACCACTTTGCC |
| Human *PCSK9* | GACACCAGCATACAGAGTGACC | GTGCCATGACTGTCACACTTGC |
| Human *BHLHE40* | TAAAGCGGAGCGAGGACAGCAA | GATGTTCGGGTAGGAGATCCTTC |
| Human *SCARB1* | GGTCCAGAACATCAGCAGGATC | GCCACATTTGCCCAGAAGTTCC |
| Human *LDLR* | GAATCTACTGGTCTGACCTGTCC | GGTCCAGTAGATGTTGCTGTGG |
| Human *GAPDH* | GTCTCCTCTGACTTCAACAGCG | ACCACCCTGTTGCTGTAGCCAA |

**Western blotting**

Protein analyses in liver were performed on ~ 25 ug total lysates. Cyro-pulverized liver tissue was incubated in RIPA buffer (Cat #89900Thermo Fisher Scientific) supplemented with 1x protease/phosphatase inhibitor (Cat #78440, Thermo Fisher Scientific) for 30 mins and sonicated for 10 secs twice. The samples were then centrifuged at 12,000 rpm for 10 mins at 4°C. Supernatant containing the protein is transferred into a new tube and used as a total lysate. For total cell lysates from culture cells, cells were harvested and resuspended in RIPA buffer containing 1x protease and phosphatase inhibitors. Lysates were incubated for 30 mins and centrifuged at 12,000 rpm for 10 mins at 4°C. The supernatant was used as a total cell lysate. The protein concentrations of the supernatants were determined using the Pierce BCA Protein Assay kit (Cat #23225, Thermo Fisher Scientific). Equal amounts of protein were separated using SDS-PAGE and transferred to a PVDF membrane (Cat #1620177, BioRad). Membranes were blocked in 5% milk in TBST for 1 hour at room temperature and then incubated overnight at 4°C with primary antibodies: PCSK9 (Cat #A7860, 1:1000, ABclonal), BHLHE40 (Cat #A6534, 1:1000, ABclonal), SR-B1 (Cat #A0827, 1:1000, ABclonal), LDLR (Cat #A14996, 1:1000, ABclonal), followed by incubation with anti-rabbit IgG, HRP-conjugated secondary antibody (Cat #7074, 1:5000, Cell Signaling) for 1 hour at room temperature. Immunoreactive bands were visualized by chemiluminescence using Pierce Enhanced Chemiluminescent western blotting substrate (Cat #32106, Thermo Fisher Scientific)

**RNA sequencing sample preparation and analysis**

RNA-seq was conducted on RNA extracted from the liver tissue of wild-type versus R24K homozygous mice. Each liver was immediately snap frozen in 1 ml TRIsure (Bioline, BIO-38033) using liquid Nitrogen. RNAs from each heart were extracted individually and re-purified using the RNeasy Mini Kit (Cat #74104, Qiagen). RNA-seq was performed at the DNA core (CCHMC). 150 ng – 300 ng of total RNA determined by Qubit (Invitrogen) high-sensitivity spectrofluorometric measurement was poly-A selected and reverse transcribed using Illumina's TruSeq stranded mRNA library preparation kit (Cat #20020595, Illumina, San Diego, CA). Each sample was fitted with one of 96 adapters containing a different 8 base molecular barcode for high level multiplexing. After 15 cycles of PCR amplification, completed libraries were sequenced on an Illumina NovaSeqTM 6000, generating 20 million or more high quality 100 base long paired end reads per sample. A quality control check on the fastq files was performed using FastQC. Upon passing basic quality metrics, the reads were trimmed to remove adapters and low-quality reads using default parameters in Trimmomatic [Version 0.33]. The trimmed reads were then mapped to mm10 reference genome using default parameters with strandness (R for single-end and RF for paired-end) option in Hisat2 [Version 2.0.5]. Next, the transcript/gene abundance was determined using kallisto [Version 0.43.1]. We first created a transcriptome index in kallisto using Ensembl cDNA sequences for the reference genome. This index was then used to quantify transcript abundance in raw counts and counts per million (CPM). Differential expression (DE genes, FDR<0.05) was quantitated through DESeq2. PCA was conducted using ClustVis. Gene ontology pathway enrichment was conducted using the Gene Ontology analysis tool.

**Chromatin immunoprecipitation sequencing**

Whole livers were cryopowdered using a liquid nitrogen-cooled RETSCH CryoMill. The cryopowdered tissue was then fixed in 10 ml of 1% paraformaldehyde (PFA) for 30 mins at room temperature with gently nutation. Fixation was quenched 1ml of 1.375 M glycine (Cat # BP381-5, Thermo Fisher Scientific) with gentle nutation for 5 min at room temperature. After centrifugation at 3000g for 5 mins at 4$^{o}$C, the pellet was resuspended in cell lysis buffer as per reported conditions, supplementing the cell lysis buffer with cytochalasin B (3 ug/ml) and rotating for 10 min at 4$^{o}$C. Nuclei were pelleted at 300g for 10 min at 4$^{o}$C and subsequently processed following the reported protocol with the adjustment of adding cytochalasin B (3ug/ml) into all solutions for chromatin preparation and sonication, antibody incubation, and wash steps. Chromatin was then sonicated for 15 cycles (30s, high power, 30s pause, and 500 µl volume) in a water bath sonicator set at 4$^{o}$C (Bioruptor 300. Diagenode, Denville, NJ). After centrifuging at 10,000g for 10 min at 4$^{o}$C, sheared chromatin was checked on agarose gel for a shear band comprised between 150 and 600 bp. Two micrograms of chromatin were kept for pooled input controls, whereas 50 ug of chromatin was used for each pull-down reaction in a final volume of 2ml rotating at

4°C overnight. Rabbit polyclonal anti-GR (Cat # A2164, 1:100, ABclonal) was used as a primary antibody. Chromatin complexes were precipitated with 100 µl of Sheep Dynabead M-280 (Cat #11204, Thermo Fisher). After washing and elution, samples were treated with proteinase K (Cat #19131, Qiagen) at 55°C, cross-linking was reversed through overnight incubation at 65°C. DNA was purified using a MinElute purification kit (Cat #28004, Qiagen) and quantified using Qubit reader and reagents. Library preparation and sequencing were conducted at the NU Genomics Core, using TrueSeq ChIP-seq library prep (with size exclusion) on 10 ng of chromatin per ChIP sample or pooled inputs and HiSeq 50-bp single-read sequencing (60 million read coverage per sample). Peak analysis was conducted using HOMER software (v4.10) after aligning fastq files to the mm10 mouse genome using bowtie2. PCA was conducted using ClustVis. Heatmaps of peak density were imaged with TreeView3. Peak tracks were imaged through WashU epigenome browser. Gene ontology pathway enrichment was conducted using the gen ontology analysis tool.

**Human iPSC cell line and maintenance**

Human iPSC line 72_3 with CRISPR knock-in for R23K in the *Nr3c1* gene locus to generate heterozygous and homozygous for GR SNP were obtained from CCHM Pluripotent Stem Cell Facility (PSCF). The hiPSCs were maintained in feeder-free conditions using mTeSR1 medium (Cat #85850, StemCell Technologies) in a humidified incubator at 37°C, 5% $CO_2$. Human iPSCs were plated on six-well plates pre-coated with Cultrex obtained from the CCHMC PSCF. The isogenic cell lines were tested and confirmed mycoplasma-free during maintenance and before differentiation process. For maintenance of hiPSC, the cells at 70% confluency were passaged using Gentle Cell Dissociation Reagent (GCDR) (Cat #100-0485, StemCell Technologies) into medium clumps. The colonies were resuspended in mTeSR™1 medium with 10 µM Y-27632 (PSCF, CCHMC) and passaged at split ratios ranging from 1:6 to 1:9 as appropriate.

**Human Albumin ELISA**

Cell supernatant containing the cell culture media from mature hiPSC-hepatocytes was collected and centrifuged at 2000 x g for 10 mins to remove debris. Centrifuged samples were diluted 1:5 in Sample Diluent NS provided in the kit (Cat # ab179887, Abcam) and assayed according to the manufacturer's instructions.

**Isolation of Primary mouse hepatocytes**

Primary hepatocytes were isolated from $GR^{ref/ref}$ (control), $GR^{ref/ALT}$ (het), and $GR^{ALT/ALT}$ (homo) mice with collagenase perfusion method. The mice were anesthetized, and the inferior vena cava (IVC) was cannulated with a 24-gauge needle. HBSS – (Cat #14175095, Thermo Fisher Scientific) containing 0.5 mM EDTA (Cat # AM9260G, Thermo Fisher Scientific) was perfused to chelate calcium. Next, HBSS + (Cat #14025092, Thermo Fisher Scientific) containing 0.3 mg/ml collagenase X (Cat #035-17861, FUJUFILM Wako Chemicals) was perfused to dissociate extracellular matrix of the liver. After the liver dissection, cells were filtered with 100 µm mesh cell strainer (Cat #08-771-19, Fisher Scientific), and the hepatocytes were purified by 40% Percoll (Cat # P1644, Sigma) gradient centrifugation method. Hepatocytes were suspended in William's E medium (Cat

#12551032, Thermo Fisher Scientific) supplemented with 10% FBS (Cat # S11150, R&D systems) and 1x Anti-Anti (Cat #15240062, Thermo Fisher Scientific) for overnight and then replaced the next day with fresh medium.

**Immunostaining and image analysis**

Cells plated on cultrex-coated dishes containing sterile cover glasses were washed gently with 1x DPBS and fixed with Fixation solution (2% formaldehyde in 1x PBS) for 15 mins at room temperature. The cells were washed 3 times with 1x DPBS and treated with permeabilization reagent (1% triton X-100 in 1x DPBS) at 37°C for 30 mins and then at room temperature for 10 mins. Next, the cells were blocked with blocking buffer (10% normal donkey serum in 1x DPBS) for 1 hour at room temperature and stained with primary antibodies: Nanog (Cat #D73G4, 1:200, Cell Signaling), OCT4 (Cat #A7920, 1:200, ABclonal), and Albumin (Cat #A1363, 1:200, ABclonal) diluted in 10% Donkey serum in 1x DPBS overnight. Next day, the cells were washed twice with 1x DPBS and stained with secondary antibodies: Alexa Fluor® 594 AffiniPure Donkey Anti-Rabbit IgG (H+L) (Cat #102649-732, 1:300, VWR), and Alexa Fluor® 488 AffiniPure Donkey Anti-Rabbit IgG (H+L) (Cat #102649-726, 1:300, VWR) diluted in 10% Donkey serum in 1x DPBS for 1 hour at room temperature. Cells were washed three times in 1x DPBS. The coverslips were mounted on slides and imaged with Nikon Eclipse Ti – U microscope.

**Nuclear and cytoplasmic protein isolation**

The separation of nuclear and cytoplasmic proteins was performed using the NE-PER Nuclear and Cytoplasmic Extraction Kit (Invitrogen #78835). Briefly, 100 mg of skeletal muscle tissue was homogenized, and 1 mL of ice-cold CERI solution was added. The sample was vortexed vigorously for 15 seconds at high speed. Following a 10-minute incubation on ice, 55 µL of ice-cold CERII solution was added, vortexed, incubated for 1 minute, and then centrifuged at 16,000g for 5 minutes. The supernatant, containing the cytoplasmic fraction, was carefully transferred to a new 1.5 mL Eppendorf tube. The remaining pellet was resuspended in 500 µL of ice-cold NER solution and placed on ice for 40 minutes, with vortexing every 10 minutes for 1 second. After the incubation, the sample was centrifuged at 16,000g for 10 minutes. The resulting supernatant, containing the nuclear fraction, was transferred to a new 1.5 mL Eppendorf tube and stored at -80°C until further analysis.

**Co-Immunoprecipitation (Co-IP) Protocol**

The Co-immunoprecipitation (Co-IP) protocol involves capturing protein complexes using an antibody specific to one of the complex members, coupling the antibody to magnetic beads, isolating and eluting the complex, and verifying the components via western blot analysis. The Universal Magnetic Co-IP Kit (Active Motif, 54002, Carlsbad, CA, USA) was used along with the appropriate antibodies (specified in Table 2) and a control IgG (2 µg) to pull down the protein complex. A total protein extract of 800 µg was prepared in a final volume of 500 µL using the complete Co-IP/Wash buffer. The extract was incubated overnight at 4°C on a rotator with the specific antibodies and IgG control. Following incubation, Protein G magnetic beads (Invitrogen #80105G) were added to the mixture and incubated at room temperature for 1 hour. The magnetic beads were then separated using a magnetic separator, and the supernatant was discarded. The beads, containing the captured protein complexes, were washed three times with IP wash buffer. For elution, the beads were resuspended in 50 µL of 2X loading dye and heated at 100°C for 5 minutes. The protein complexes were then separated from the beads using a magnetic stand, and the eluted proteins were transferred to a new tube. Finally, the samples were loaded onto a 10% SDS-PAGE gel, with 20 μL of each sample per lane, for further analysis.
